# Single-cell Gene Regulation Network Inference by Large-scale Data Integration

**DOI:** 10.1101/2022.02.19.481131

**Authors:** Xin Dong, Ke Tang, Yunfan Xu, Hailin Wei, Tong Han, Chenfei Wang

**Affiliations:** Department of Urology, Tongji Hospital, Frontier Science Center for Stem Cells, School of Life Sciences and Technology, Tongji University, Shanghai 200092, China

**Keywords:** gene regulation network, scATAC-seq, transcription factor, ChIP-seq, data integration

## Abstract

Single-cell ATAC-seq (scATAC-seq) has proven to be a state-of-art approach to investigating gene regulation at the single-cell level. However, existing methods cannot precisely uncover cell-type-specific binding of transcription regulators (TRs) and construct gene regulation networks (GRNs) in single-cell. ChIP-seq has been widely used to profile TR binding sites in the past decades. Here, we developed SCRIP, an integrative method to infer single-cell TR activity and targets based on the integration of scATAC-seq and a large-scale TR ChIP-seq reference. Our method showed improved performance in evaluating TR binding activity compared to the existing motif-based methods and reached a higher consistency with matched TR expressions. Besides, our method enables identifying TR target genes as well as building GRNs at the single-cell resolution based on a regulatory potential model. We demonstrate SCRIP’s utility in accurate cell-type clustering, lineage tracing, and inferring cell-type-specific GRNs in multiple biological systems. SCRIP is freely available at https://github.com/wanglabtongji/SCRIP.

## Introduction

Gene regulation is the fundamental basis of many biological processes, including development and differentiation, disease occurrence, and progression. Recently, many single-cell technologies have been developed to investigate gene regulation mechanisms from diverse genomic aspects, such as transcriptomes(1), epigenomes(2), or 3D structures(3). Among them, the single-cell sequencing assay for transposase-accessible chromatin (scATAC-seq) has enabled the profiling of the genome-wide chromatin accessibility landscapes in single cells (4). The most powerful application of the scATAC-seq data is to understand how specific transcription regulators (TR), including transcription factors (TF) and chromatin regulators (CR), bind to the genome and regulate their target genes. Constructing the gene regulatory networks (GRNs) is crucial for understanding the roles of different TRs in regulating development trajectories and disease traits.

Although scATAC-seq has been widely used to tackle gene regulation and their association with phenotypes, several questions remain unsolved. First, the chromatin accessibility captured by scATAC-seq only reflects the overall regulatory potential and cannot identify the binding of exact TRs. Existing methods like chromVAR(5), scFAN(6), and SCENIC(7) integrate sequence features like motifs to evaluate TF activity in each cell. However, motif-based methods cannot discriminate factors of the same TF family that have similar motifs, and also failed to evaluate factors with indirect DNA binding such as CRs. Second, the scATAC-seq data is very sparse and noisy as only two strands of DNA can be captured within a cell. Methods like Signac(8), EpiScanpy(9), MAESTRO(10), and SCALE(11) enhanced the signals by using different latent features, however, the algorithm-defined features were mostly analyzed at the cell type level and cannot be directly linked to the single-cell TR activity. Last but most important, none of these methods can identify the TR targets in each cell, and constructing the GRNs at the singlecell level is still not feasible with scATAC-seq data alone. Therefore, new methods with the potential to address TR binding enrichment and identify its associated target genes at the single-cell level are highly needed for scATAC-seq data.

Chromatin Immunoprecipitation Sequencing (ChIP-seq) (12, 13) is a direct way to uncover TRs binding in the genome and determine their target genes at the bulk cell level. Compared to motifs, ChIP-seq is more accurate in defining cell-type-specific TR binding sites and investigating the genomic distribution for many non-DNA-binding CRs. In the past decades, numerous TR ChIP-seq data have been generated for different cell lines, tissues, and species (14–16). Several projects, such as CistromeDB (16), ENCODE (15), and Epigenome Roadmap (17) have curated a large collection of high-quality TR ChIP-seq data. Integrating the large-scale ChIP-seq dataset with the motif information will definitely improve the prediction of TR enrichment in the scATAC-seq data. However, several issues need to be addressed before integration. First, the large-scale ChIP-seq reference should be uniformly processed with standard quality control metrics to remove the potential low-quality data. Besides, while the antibody-affinity and the signal-to-noise ratio might be diverse for different TRs, the enrichment based on TR ChIP-seq peaks should be carefully adjusted and normalized. Finally, an efficient interval searching algorithm is needed for identifying enriched TRs from a large-scale genomewide TR reference (18).

The major obstacle to investigating single-cell GRNs is the lack of single-cell ChIP-seq data. While several recently developed techniques such as scCUT&RUN(19), scCUT&Tag(20), and scCUT&Tag-pro(21) could successfully generate ChIP-seq profiles at the single-cell level, however, most of the data were generated for high abundant histone modifications (HMs) rather than TRs. Although several attempts have been performed on specific TRs(22), they are highly dependent on the quality of the TR antibodies and usually have extremely fuzzy signals at the single-cell level. Regulatory Potential (RP) models have been widely used to identify TR targets for bulk ChIP-seq samples (23–25). The integration of TR ChIP-seq and scATAC-seq data has the potential for evaluating the single-cell TR binding site using the RP model, which could be based on the imputed TR ChIP-seq peaks at the single-cell level.

Here, we present a computational method SCRIP, which integrates a large-scale TR and motif reference for evaluating TR activity as well as constructing single-cell GRNs based on scATAC-seq data. SCRIP includes a high-quality TR reference covering 1,252 human TRs and 997 mouse TRs. Based on this large-scale reference, SCRIP showed superior performance in evaluating single-cell TR activity and performing TR-based clustering and lineage tracing analyses. In addition, SCRIP could accurately reconstruct the single-cell GRNs based on imputed ChIP-seq peaks at the single-cell level. We demonstrated the usability of SCRIP on multiple biological systems including peripheral blood mononuclear cell (PBMC), hematopoietic stem cell (HSC) differentiation, human organ development, and basal cell carcinomas (BCC).

## Materials and Methods

### Data collection and generation of SCRIP index

We downloaded the uniformly processed ChIP-seq datasets from the Cistrome Data Browser (16, 26, 27) through the “batch download” function. In total, we obtained bed files of 11,348 human and 9,060 mouse TR ChIP-seq datasets, and 11,079 human and 10,944 mouse HM ChIP-seq datasets. Since the auto-parsed metadata included many mistakes, we systematically curated the annotation of factors, cell types, and tissues. To ensure the quality of the datasets, we used the following criteria to filter the TR datasets: the raw sequence median quality score was greater than 25, the percent of uniquely mapped reads was greater than 50%, PBC (PCR bottleneck coefficient) was greater than 0.8, the number of fold 10 peaks was greater than 100, FRiP (Fraction of Reads in Peaks) was greater than 0.01 and the number of top 5,000 peaks overlapping with union DHS is greater than 70%. To acquire the high confidence peaks, we only kept the 5-fold enrichment peaks in each peak set. Then, we removed the datasets with less than 1000 peaks. After filtering, we obtained 2,314 human and 1,920 mouse TR ChIP-seq datasets, covering 671 and 440 TRs respectively (Fig. S1). We also filtered histone modification datasets with the same criteria as above. According to previous reports, active histone modifications such as H3K4me1/2/3 or H3K27ac tends to be presented in the open chromatin region, while repressive histone modifications such as H3K27me3 or H3K9me3 are enriched at the heterochromatin region and have very few overlaps with the scATAC-seq peaks (28–30). Thus, we only retained the histone modifications with active functions, including H3K4me1/2/3, H3K9ac, and H3K27ac. We obtained 1,678 human and 1,013 mouse HM ChIP-seq datasets covering 5 HMs (Fig. S2).

To improve the coverage of transcription factors, we downloaded the motif information, including 7,704 human and 7,000 mouse TF PWMs (Position Weight Matrix), from the Cis-BP database (31). We combined PWMs from the same TF and converted the format to the HOMER (32) format. Then we scanned the motifs on the hg38 or mm10 genome with the HOMER and obtained the genome intervals where motifs appeared. We overlapped the scanning intervals with the ENCODE ccRE (candidate cis-regulatory elements) (33) list and Cistrome union DHS (DNase-I hypersensitive sites) list and removed intervals with the intersection of the blacklist. To make the motif sites comparable to the ChIP-seq dataset, we extended the length of each scanned motif site to 340bp, which is the average length of ChIP-seq peaks. For those motifs that have much more binding sites than others, we only kept the top 25k binding sites, which were the average of filtered peaks in ChIP-seq, by filtering out the low confidence motif sites using p-values. In total, we obtained 916 human and 816 mouse motif-scanned pseudo peaks (Fig. S1e, f).

Next, we combined the ChIP-seq peak sets and motif-scanned pseudo peak sets as reference datasets to build the search index. To calculate the similarity between reference datasets and the scATAC-seq datasets, we introduced GIGGLE (18), a fast genomics search engine, into SCRIP. We sorted the bed files, compressed them into gz format, and built the index with GIGGLE. In addition, we also included the peaks number, metadata of the datasets, and original bed files in the index. Overall, the human TR index covers 1,252 TRs and the mouse covered 997 TRs (Fig. S1e, f).

### TR activity score calculation

#### Normalization for removing biases

The SCRIP takes the scATAC-seq peak by count matrix or bin count matrix as input. For the sake of getting the comparable TR activity of each reference dataset in each cell, SCRIP first calculates the number of peak overlaps between each cell and the ChIP-seq peaks set or motif-scanned intervals set by GIGGLE. SCRIP records the number of overlap peaks to build the matrix *M*, where the column is cell *i* and the row is dataset *j*, and the content is the number of overlapped peaks. To remove the bias from the peak number of the datasets and the total length of single-cell peaks, SCRIP normalizes the matrix by:

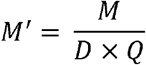

where *D* is a *j* length vector that records the number of peaks of each dataset, and *Q* is an *i* length vector that records the number of base-pair in each cell per 100 million. Then the *M*’ matrix is the normalized peak overlap matrix, which scores the relative enrichment of TRs in different cells. *M*’ is further normalized by the average score of each TR dataset to scale the enrichment for different TRs:

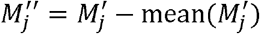

#### Deduplication for redundant TR datasets

As *M*” still contains duplicate ChIP-seq datasets or motifs for the same TR, we remove the duplicate datasets and only keep the column with the largest score for the same TR *k*:

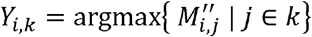

where the *Y_i,k_* is the TR enrichment score matrix. In this step, the best-matched dataset of each cell is determined independently according to the largest score within each cell. Here, we set this maximum strategy as default in SCRIP due to its superior performance (Fig. S3), but also provide an average strategy option, which uses the average of all same TR datasets to represent the TR enrichment score.

#### Scaling the TR enrichment scores

Next, to stabilize the TR enrichment score and compress the outliers, we introduce the logistic sigmoid function to each TR. The z-score is a step of the sigmoid function, which will shift the data range around 0 for effective logistic sigmoid transformation:

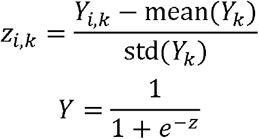

Finally, SCRIP applies z-score normalization to each cell to ensure that the TR activity scores after sigmoid normalization has a similar dynamic range and achieve better clustering performance:

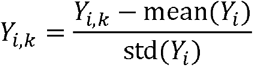

To better understand the TR enrichment score calculation and normalization, the distribution changes along with score calculation and normalization were shown in Fig. S4.

#### TR targets modeling

To acquire scaled and dependable targets of a TR, SCRIP first imputes the potential binding sites of each TR in each cell. For a specific TR, the best match ChIP-seq dataset of each cell was determined in the TR activity score calculation step. SCRIP then imputes the potential TR binding sites by overlapping the ChIP-seq peak sets with scATAC-seq peaks or intervals of each cell. Since some TR ChIP-seq datasets do not have a sufficient number of peaks and ChIP-seq peak sets are performed on bulk tissue, which may include the peaks from other celltype, SCRIP further provides a function that uses all the best match TR peak sets of this data found to include other potential binding sites.

With the TR potential binding sites, we can measure the effect on other genes of this TR, or determine the target of this TR, with the RP score model. The RP score of a gene is its likelihood of being regulated by a TR and has been used in several previous studies (10, 23–25). In general, the RP models could be classified as signal RP models and peak RP models, which are based on scaled signals or binarized peaks, separately. Compared to the signal RP model, the peak RP model used in Cistrome-GO (25) and MAESTRO (10) is more compatible with the binarized signal of scATAC-seq and thus is introduced here. In SCRIP, the formula of the RP score calculation is shown below:

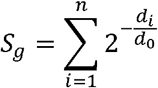

where the d_0_ is the half-decay distance or regulatory range, and it can be changed by users. The n denotes the number of binding sites near the TSS of gene g. To save the computation time, SCRIP only takes genes within 15 d0 into account as the score will be less than 0.0005 if a peak is over 15d_0_. The d_i_ is the distance between the i_th_ peak’s center and TSS. This is defined as the simple RP model in SCRIP. The enhanced RP model takes the exon information and nearby genes into account. If a peak is located at the exons region of a gene, the score of this peak is set to 1 and further normalized by the total exon length of the gene. If a peak is located in the promoter or exon regions of any nearby genes, then the score of this peak is set to 0. Also, for specific TRs, we added a “auto” mode to automatically determine the d_0_ by the percentage of the TR peaks on promoter regions (1kb around TSS). TRs with more than 20% peaks on promoter regions are defined as promoter-type TRs and use 1k as the half-decay distance. Other TRs are defined as enhancer-type TRs and use 10k as the half-decay distance. We compared the performance of the simple RP model and enhanced RP model, which use different half-decay distances for different types of TRs (1k for promoter-type TRs, and 10k for enhancer-type TRs). We applied different RP models to identify the targets for 140 TRs from the PBMC scATAC-seq dataset. To evaluate the performance, we calculated the expression correlation between TR and its top 500 targets using matched PBMC scRNA-seq and used the number of true targets (abs correlation >= 0.3 and p-value <= 0.01) as the evaluation metric. We first compared the performance of using different regulatory ranges. 1k group represents the half-decay for all TRs were set as 1k, similar for the 10k group. For the auto group, the half-decay is determined by the percentage of the TR peaks on promoter regions. Clearly, the auto half-decay strategy shows an overall higher number of true targets compared to 1k and 10k strategies (Fig. S5a). We also compared the performance of the simple versus enhanced RP model. Interestingly, the simple RP model seem to have slightly better performance than the enhancer RP model considering all TRs (Fig. S5b). However, for factors like IRF8 and CEBPA, the enhanced RP model shows better performance, while TBX20 and EBF1 have a different trend. Besides, the two models share a great number of target genes for the same factor (Fig. S5c-d). These results suggest that the simple and enhanced models do not show a significant difference in identifying target genes. We set the simple RP model and auto half-decay distance as the default parameter for SCRIP, but also provided the enhanced model for users to choose from. With the RP score, we can rank the target genes for each TR, and obtain the top-ranked target genes to build the gene regulatory network.

### Data processing on different scATAC-seq datasets

#### PBMC multiome dataset

##### Preprocessing and TRs activity

The PBMC multiome data was downloaded on the 10X genomics website. The peaks were called using Cell Ranger ATAC 2.0 by fitting the peak signals using a Zero-Inflated Negative Binomial model, combining scATAC-seq reads as a bulk sample. In the scRNA-seq data, cells with less than 200 genes and genes with less than 3 cells were removed. We only retained the cells with both RNA-seq counts and ATAC-seq counts. We clustered the scRNA-seq data with the Louvain clustering algorithm and annotated the cell type by cell markers (Fig. S6a, c, d). Then, we transferred the cell type labels to scATAC-seq data by the matched cell barcodes (Fig. S6b, e). We applied the SCRIP to the filtered scATAC-seq peak count matrix with the default parameters to evaluate the activity of TRs. The activity scores of different TRs were used to draw heatmap with clustermap of seaborn and project to UMAP with scanpy (Fig. 2a, S7).

**Figure 1.**
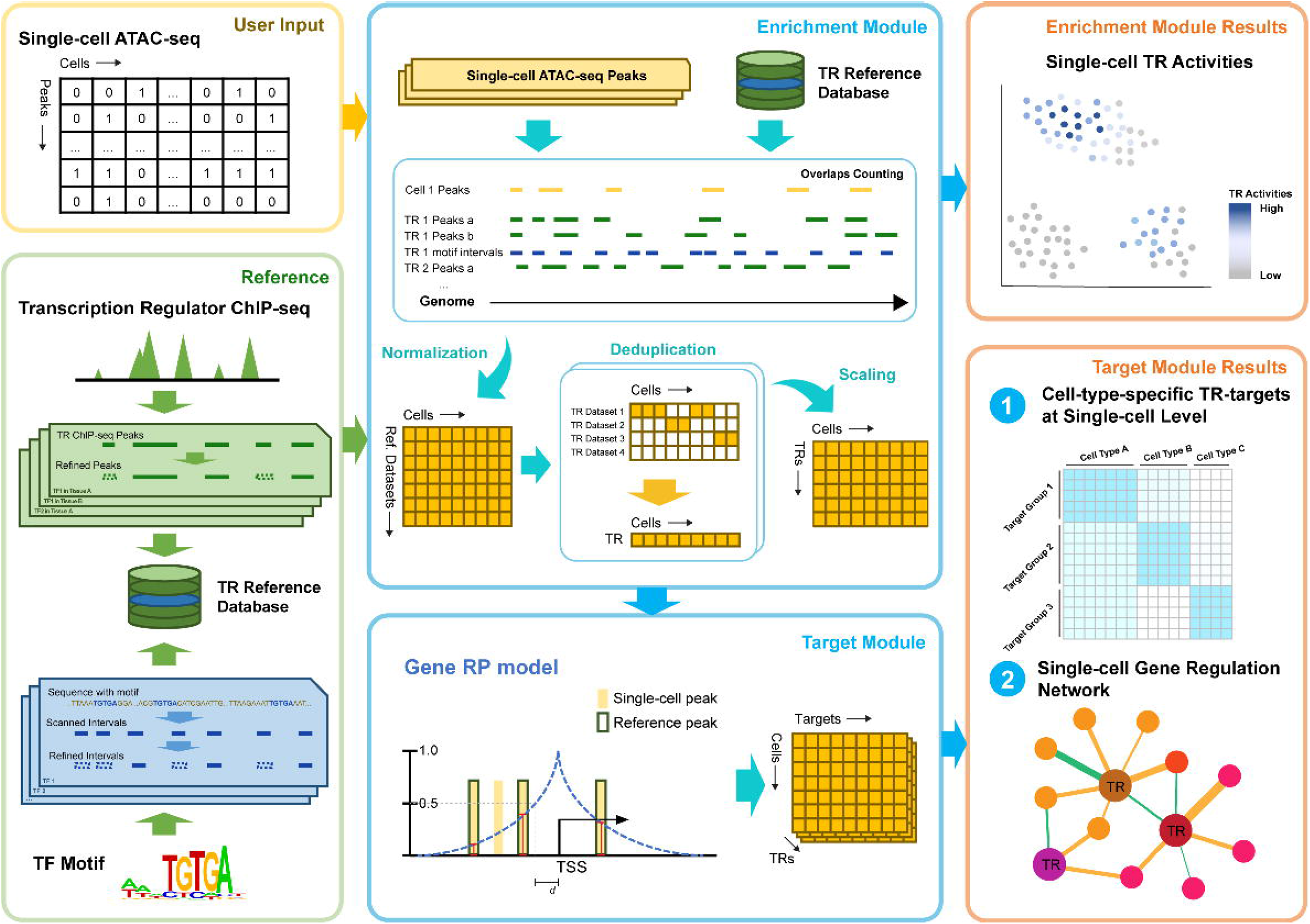
Workflow of SCRIP. Schematic of SCRIP workflow. SCRIP takes the feature count matrix of scATAC-seq as input. The TR ChIP-seq and motif reference datasets were built based on Cistrome ChIP-seq data and Cis-BP motifs, with careful curation. For the Enrichment Module, the overlaps of scATAC-seq and reference datasets are firstly counted using GIGGLE and further normalized. Then the scores for the same TR were merged and only kept the score of the best-matched dataset for every TR in every cell. Finally, the TR scores were scaled and output. For the Target Module, the best-matched ChIP-seq peaks are combined with scATAC-seq peaks to determine the target gene using the RP model. SCRIP outputs the TR activity, differential targets between cell types, and GRNs at the single-cell level.

**Figure 2.**
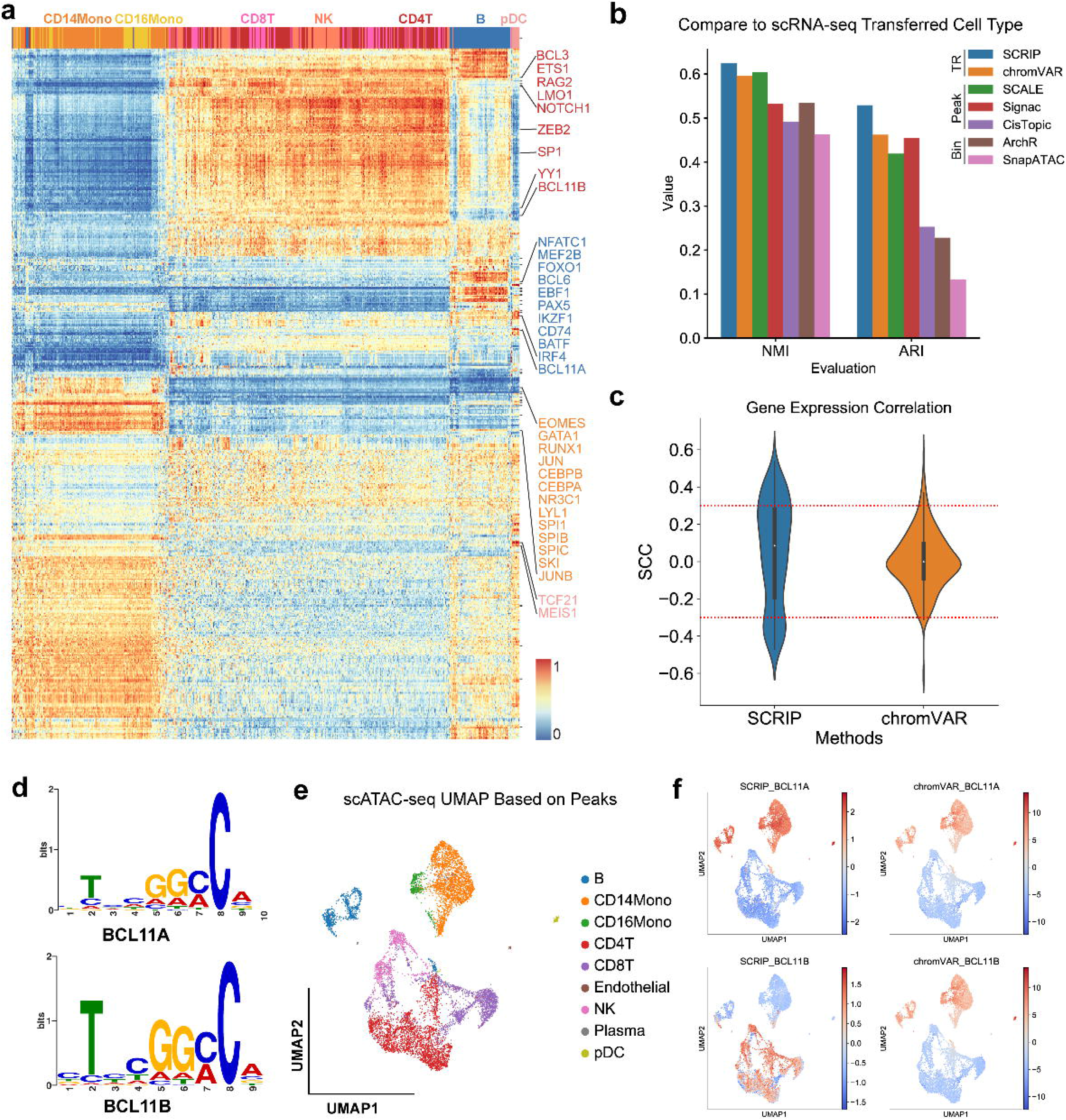
SCRIP achieves better performance on the PBMC dataset. **a.** Heatmap of TR activity with different cell types. The X-axis denotes the unsupervised clustering results with the TR activity score. **b.** The clustering performance among SCRIP, chromVAR, SCALE, Signac, CisTopic, ArchR, and SnapATAC. Y-axis: NMI scores or ARI scores. **c.** Distribution of Spearman’s correlation coefficients of TR activity and TR expression from SCRIP and chromVAR. **d.** The SeqLogo of BCL11A and BCL11B motifs. **e.** Annotation of scATAC-seq with scRNA-seq transferred labels. UMAP was generated by the peak count matrix. **f.** SCRIP and chromVAR TR activity results of BCL11A and BCL11B in PBMC dataset.

##### Clustering performance comparison

We compared the clustering performance between TR-based tools (SCRIP, chromVAR), peak-based tools (SCALE, Signac, CisTopic), and bin-based tools (ArchR and SnapATAC) in the PBMC dataset. The scATAC-seq data was preprocessed into TR-cell, peak-cell, or bin-cell matrix to meet the requirements of each tool. All tools were applied to scATAC-seq data with their default parameters. In addition, we use the chromVAR motifs as the motif reference in the process of chromVAR. The Louvain algorithm in Seurat or scanpy was used to perform clustering with the tools. Normalized mutual information score (NMI) and Adjusted rand index (ARI) were calculated with the python package sklearn (Fig. 2b).

##### TR activity and expression correlation

SCRIP TR activity score and chromVAR z-score were used to calculate the Spearman correlation coefficients (SCC) with gene expression. Only the 468 TRs that appear in SCRIP, chromVAR, and gene expression were used to calculate the correlation (Fig. 2c, S8). We also separated the positive and negative correlations and compared them individually. We defined the TRs with SCC > 0.3 with their expression, and the p-value < 0.01 as high-confidence positive regulators, and with SCC < −0.3 and the p-value < 0.01 as high-confidence negative regulators. We count the number of high-confidence regulators for both positive and negative regulators between SCRIP and chromVAR respectively.

##### Datasets selection evaluation

We evaluated SCRIP’s ability for identifying the best-matched dataset using POLR2A, a TR that has the most abundant ChIP-seq references (Fig. S1c-d, >150 high-quality datasets). We count the number of identified TR datasets for each cell type and the number of cells for specific TR datasets (Fig. S9). Considering that not all of the single-cells, especially for the rare population such as mast cells, could find TR datasets with matched cell types. We did further analyses to compare the performance of using cell-type-matched strategy, average strategy (average the score for the same TR), and maximum strategy (highest score within the same TR, current model in SCRIP). Only 340 TRs were used for the cell-type-matched strategy due to the relatively low cell type coverage of many TRs. Finally, we checked the TR enrichment distributions of several cell-type-specific TRs using different strategies (Fig. S3).

##### H3K27ac targets determination

To capture the loci of rare population cell types, we convert the 10X scATAC-seq data and scCUT&Tag-pro data to the bin-cell matrix with 500 bp. We applied the SCRIP impute function to the scATAC-seq data with HM reference to impute the loci of H3K27ac modifications. The genome track was built by merging the same cell type and normalizing by min-max normalization (Fig. 3b, S10). With the imputed H3K27ac bin-count matrix, we applied the SCRIP target function to calculate the RP score of each gene and determine the key affected genes with the H3K27ac modification. We applied the same algorithm to calculate the RP score of the scCUT&Tag-pro dataset and the original scATAC-seq dataset. We merged the T cells and monocytes with the maximum RP score, which reflects the driven peaks in the cell type. Also, a bulk T cell and a monocyte dataset’s RP score were obtained from Cistrome DB. Then, we calculated the Spearman correlation with the scCUT&Tag-pro dataset and showed the overlap of target genes between the top 1,000 RP score targets of each dataset (Fig. 3c).

**Figure 3.**
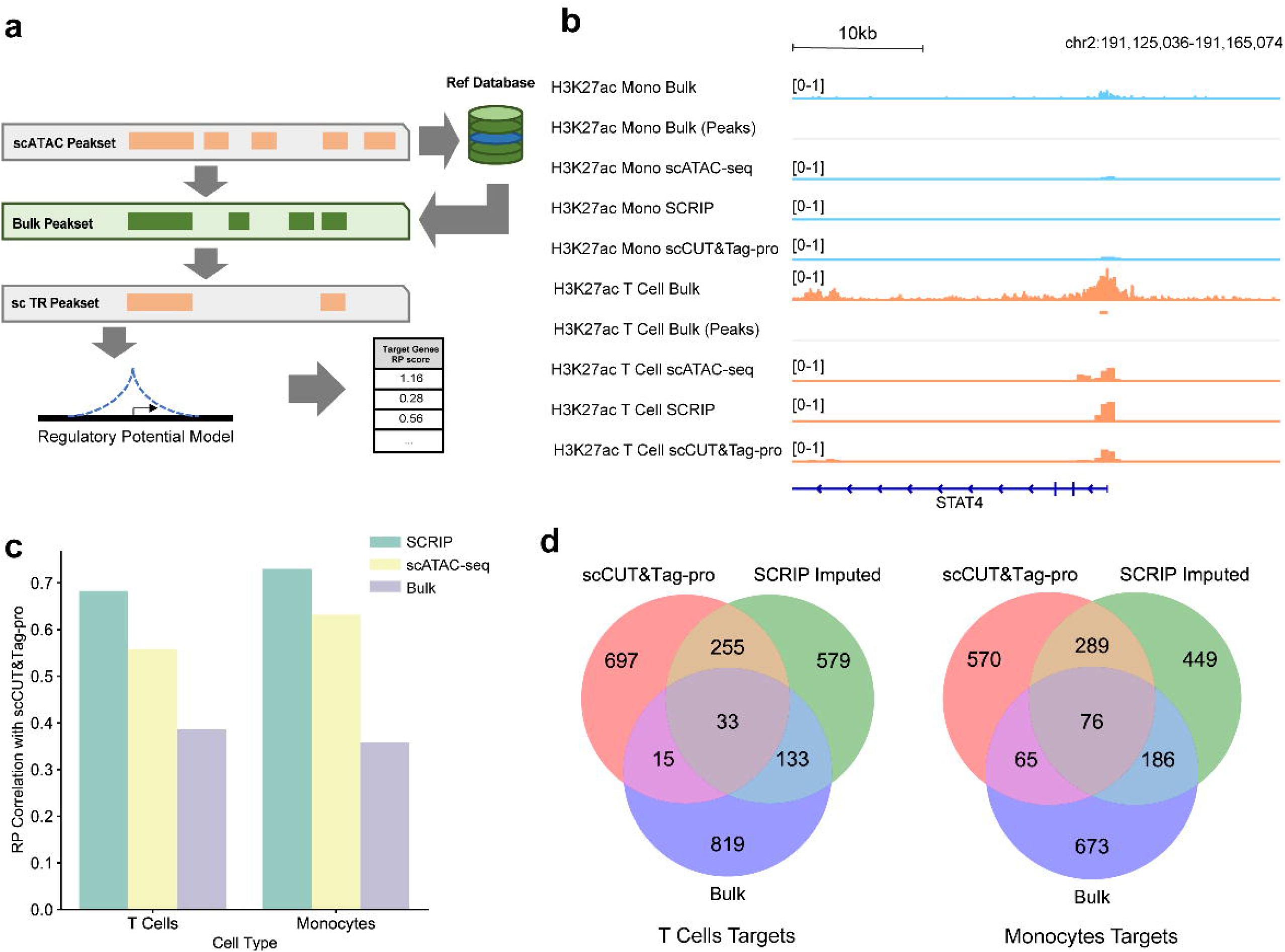
SCRIP enables finding targets from the scATAC-seq data. **a.** A simple schematic of SCRIP workflow on determining target genes. **b.** Genome track of monocytes and T cells on H3K27ac signals at STAT4. Light blue: Monocytes; Orange: T cells. Bulk tracks are read level; scCUT&Tag-pro and SCRIP-inferred tracks are 500 bp bin level. In single-cell tracks, the height of the signal denotes the normalized cell number. **c.** Spearman correlation coefficients of RP scores between SCRIP imputed, original scATAC-seq and bulk dataset with scCUT&Tag-pro in T cells and monocytes. **d.** Top 1000 target genes overlap among H3K27ac scCUT&Tag-pro, SCRIP imputed, and bulk dataset in T cells and monocytes.

#### HSC Dataset

##### Preprocessing

The HSC scATAC-seq peak count matrix was obtained from the GEO. LiftOver was used to convert genome build from hg19 to hg38. The HSC peaks were generated by MACS2 (34) using bulk hematopoietic data from the same study. We annotated the cell type with the labels from the original study. To evaluate the activity of TRs in each cell, we applied the SCRIP enrich function to the peak count matrix with the default parameters. We performed the unsupervised clustering with the TR activity score and calculated the NMI and ARI with the cell type annotations (Fig. S11e, Table S1). The TRs’ activity score matrix was used to do the following analysis.

##### Trajectory analysis

We applied the R packages destiny (35) to perform the trajectory analysis of HSC. To meet the requirement of data distribution of the destiny, we did an extra normalization step that centralized the activity score at the TR level after the deduplication. The top 600 most variable TRs were used to reconstruct the differential path. The k was set to 4 for the k-Nearest Neighbor (KNN) algorithm in Destiny. The tip was set to 1 to calculate the Diffusion Pseudo Time (DPT). The first two diffusion components were used to draw the diffusion map. We evaluate performances of trajectory between SCRIP, chromVAR, and peak count with the relative distance between starting cell types (HSC) and terminal cell types (monocytes, MEP, CLP, and pDC). The start position was indicated using the HSC’s average coordinates. We carry out the 0-1 normalization for the four terminal cell types using the start position and every terminal cell type cell. The cell type position was then determined by averaging the coordinates of its cells. Between the start position and cell type position, the Euclidean distance was determined. We projected the HOXA9, GATA1, CEBPB, TCF4, and other TRs activity scores to the diffusion map to show the TRs activity on each branch (Fig. 4b, d, Fig. S11f-j). To visualize the dynamic changes in the TRs’ activity in the differential lineages, we showed the TR’s activity score with the cell’s DPT (Fig. S11k-z).

**Figure 4.**
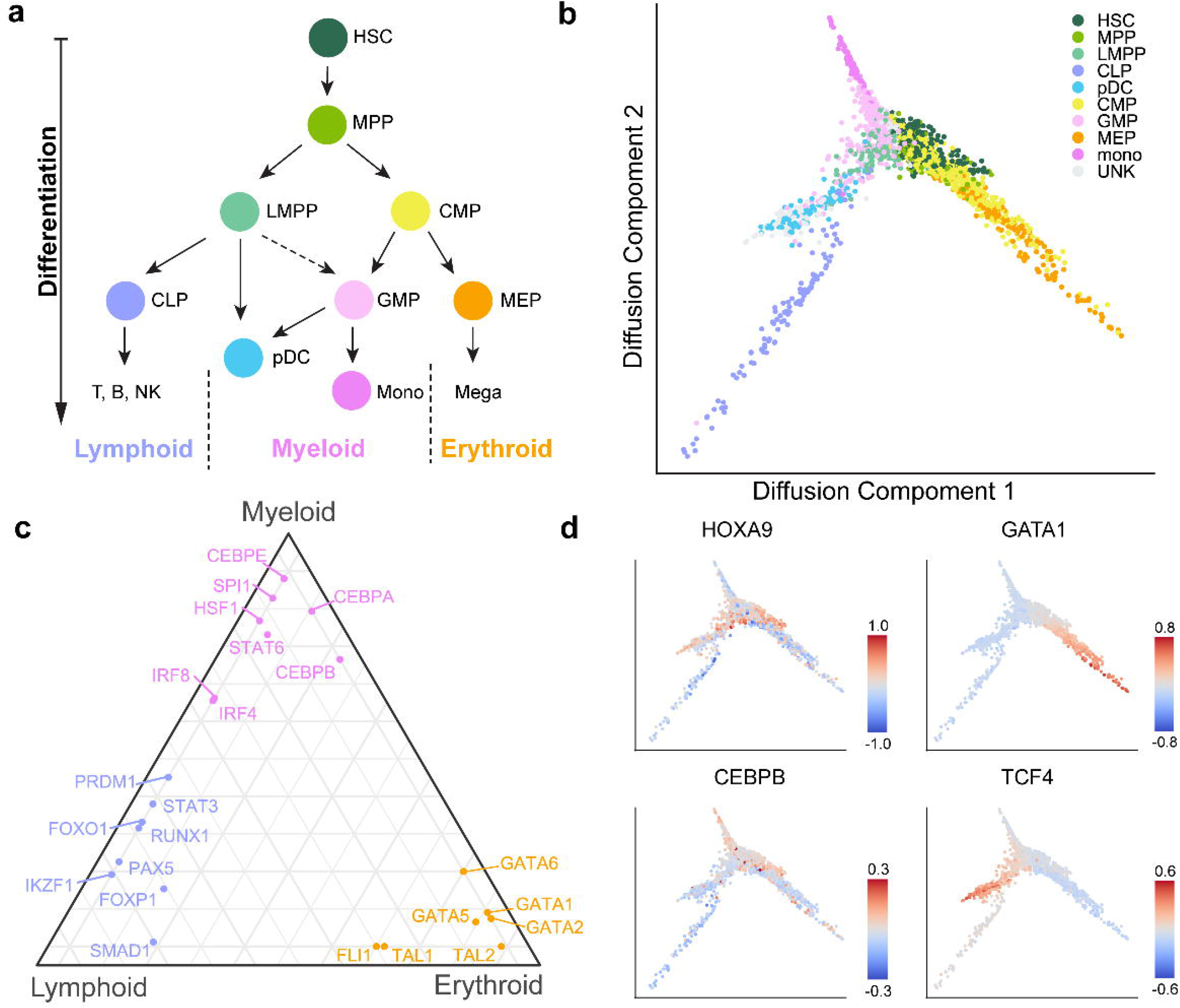
SCRIP reconstructs the path of differentiation of HSCs based on the TR activity. **a.** Schematic of HSC differentiation. **b.** Diffusion map of HSC with the cell-type annotations. MPP: multipotent progenitor; LMPP: Lymphomyeloid-Primed Progenitor; CMP: common myeloid progenitor; GMP: granulocyte-macrophage progenitors; UNK: unknown (original study annotation). **c.** Triangle plot of TRs that regulate HSC differentiation towards three main lineages. **d.** Projecting HOXA9, GATA1, CEBPB, and TCF4 activity onto the diffusion map.

##### Triangle Plot

The activity score of each TR of each cell type was calculated by averaging the TR’s activity of all cells in the same cell type. The quantile of the TR in each cell type among all cell types was used to suggest TR’s preference for each cell type. The TRs’ activity on different lineages were represented by the TR activity score of the terminally differentiated cell type. For example, we used the CLP (Common Lymphoid Progenitor) to represent the lymphoid branch, the monocytes to represent the myeloid branch, and the MEP (Megakaryocyte-Erythroid Progenitor) to represent the erythroid branch. We displayed the positions of TRs on each branch with the ggtern package (Fig. 4c).

#### Human Fetal Organ Datasets

##### Preprocessing and Clustering

The different human organs scATAC-seq datasets were obtained from the GEO, which provided the filtered peak count matrix and cell labels with the Seurat object format. The peaks of each sample were called using MACS2 by combining scATAC-seq reads as a bulk sample, then the peaks from different samples/organs were merged to generate a union peak set. LiftOver was used to convert genome build from hg19 to hg38. We applied the SCRIP enrich function to the provided scATAC-seq peak count matrix with the default parameters. Then, we used the most variable TRs in each organ and clustered them with R packages ggtree and ComplexHeatmap (Fig. 5b-d, Fig. S12a-c, Table S2). We performed the unsupervised clustering with the TR activity score and calculated the NMI with the cell type annotations. (Fig. 5a, Table S1)

**Figure 5.**
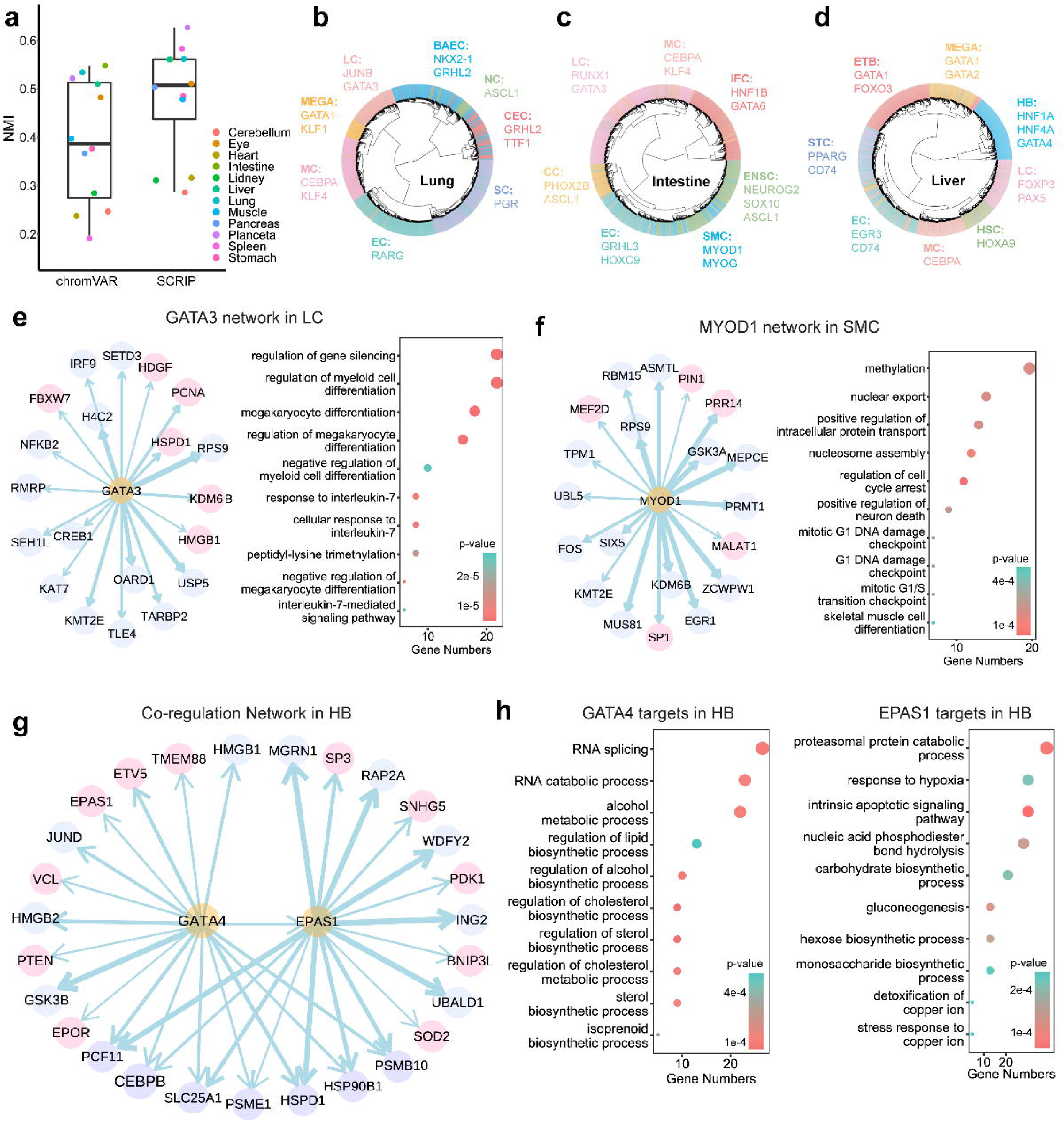
SCRIP finds the key regulators and builds GRNs in human fetal organs. **a.** Clustering performance of 12 different organs. Only NMI plots here. ARI can be checked in Table S1. **b-d.** Clustering of the human lung (b), intestine (c), and liver (d) scATAC-seq data based on TR activity. Marked TRs of specific cell types are identified by SCRIP with literature support. LC: Lymphoid cells; MEGA: Megakaryocytes; MC: Myeloid cells; EC: endothelial cells; SC: Stromal cells; CEC: Ciliated epithelial cells; NC: Neuroendocrine cells; BAEC: Bronchiolar and alveolar epithelial cells; ETB: Erythroblasts; HB: Hepatoblasts; HSC: Hematopoietic stem cells; STC: Stellate cells; IEC: Intestinal epithelial cells; ENSC: Enteric neural stem cells; SMC: Smooth muscle cells. **e.** (left) Inferred GATA3 GRN with human lung’s LC scATAC-seq dataset. Pink circles denote the target genes that are supported by previous studies. (right) GO results showed the terms enriched of GATA3 target genes. **f.** Inferred MYOD1 GRN with human intestine’s SMC scATAC-seq dataset. Pink circles denote the target genes that are supported by previous studies. (right) GO results showed the terms enriched of MYOD1 target genes. **g.** Inferred GATA4 and EPAS1 co-regulation network with HB cells of human liver scATAC-seq dataset. Pink circles denote the target genes that are supported by previous studies. **h.** GO results showed the terms enriched of GATA4 target genes and EPAS1 target genes in HB.

##### Target analysis

We applied the SCRIP impute and target functions with default parameters to determine the GATA3 target genes in the lung, the MYOD1 target genes in the intestine, and the GATA4 and EPAS1 target genes in the liver. To build the credible GRN of the four TRs, we retained the 500 cells with the highest RP in each cell type. The GRNs were built by the R package ggraph. To know the functions of TR’s targets, we selected the top 1000 target genes according to the RP score to do the gene ontology (GO) enrichment analysis with the R packages ClusterProfiler (Fig. 5e-h, Table S3).

#### BCC tumor microenvironment Datasets

##### Preprocessing

The scATAC-seq dataset of tumor cells and T cells in BCC was obtained from GEO. Cells were first clustered using a 2.5kb bin-based method, then the cells from each cluster (cell type) were merged as a pseudo bulk and the peaks were called using MACS2. LiftOver was used to convert genome build from hg19 to hg38. We applied the SCRIP enrich function to the provided scATAC-seq peak count matrix with the default parameters. The averages of TR activity in each cell type were used to plot the heatmap (Fig. 6a). The pseudo-time analysis was conducted by the custom scripts from the original study of this dataset. The UMAPs and violin plots of gene expression were obtained from the TISCH database (36).

**Figure 6.**
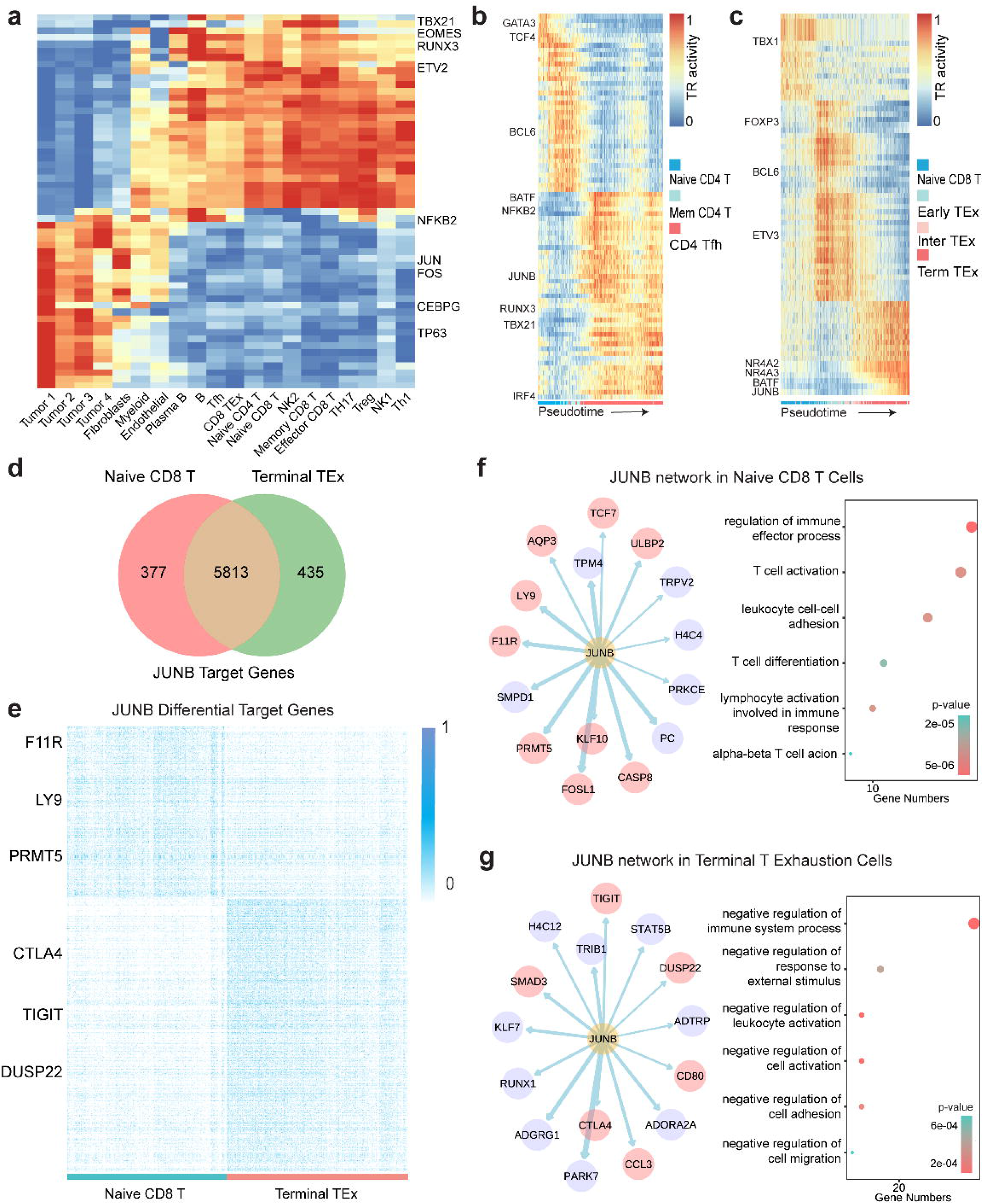
SCRIP uncovers differential targets in the tumor microenvironment. **a.** Heatmap of cell-type-specific TRs in BCC tumor microenvironment. **b.** TR activity change during pseudo-time from naive CD4 T cells to CD4 Tfh cells. **c.** TR activity change during pseudo-time from naive CD8 T cells to terminal TEx cells. **d.** Overlap of specific target genes of JUNB of naive CD8 T cells and terminal TEx cells. **e.** Heatmap of RP scores of specific target genes of JUNB of naive CD8 T cells and terminal TEx cells. RP scores were normalized to 0 to 1 with min-max normalization. **f-g.** (left) Inferred JUNB GRN in naive CD8T cells or terminal TEx cells. Pink circles denote the target genes that are supported by previous studies. (right) GO results showed the terms enriched of JUNB target genes in naive CD8T cells or terminal TEx cells.

##### Target analysis

We applied the SCRIP impute and target functions with default parameters to determine the BATF target genes in terminal TEx cells and the IRF4 target genes in CD4 Tfh cells (Fig. S13a, b). The JUNB target analysis was done on both naïve CD8 T cells and terminal TEx cells (Fig. 6f, g, Table S3). Genes with low RP scores, which are considered to have few peaks of this TR, were removed. We normalized RP with the natural logarithm and scaled it for each cell. The FindMarkers function in the R package Seurat was applied to identify the differential target genes of naïve CD8 T cells and terminal TEx cells according to the normalized RP score. Different target genes were obtained by 0.25 log fold change and 0.01 p-value (Fig. S13d). The ClusterProfiler was used for GO analysis of these target genes. The GRNs were built using the R package ggraph.

## Results

### Workflow of the SCRIP

The SCRIP workflow takes the peak count or bin count matrix of scATAC-seq data as input and outputs the TR activity score and their target genes in each cell (Fig. 1). We first built a comprehensive reference dataset to help evaluate the TR enrichment in the cells from scATAC-seq data (Fig. S1-S2). The reference dataset includes two components. The first one is a TR ChIP-seq reference based on a large collection of 20k ChIP-seq datasets from the Cistrome Data Collection (11k human TRs and 9k mouse TRs) (16, 26). We have carefully curated metadata such as factor information, tissue types, and cell types (Fig. S1a-b). Then we filtered out the ChIP-seq datasets of bad quality and removed low confidence peaks from retained datasets to generate high confidence TR peak sets covering 671 human TRs and 440 mouse TRs (Fig. S1c-d). Considering there are also TRs without ChIP-seq datasets, we also scanned motifs on the whole genome and obtained the refined intervals with high confidence. These two references were combined to generate the TR reference database containing 1,252 human TRs and 997 mouse TRs in different tissues (Fig. S1e-f).

Next, we evaluated the TR enrichment in each cell by modeling the peak overlaps between scATAC-seq peaks and TR reference. While scATAC-seq peaks are usually sparse and noisy, we first implemented an imputation step using nearest neighbor cells. Then, we calculated the intersections of each TR dataset or motifs in every single cell. This score was further normalized by the number of reference peaks and length of scATAC-seq peaks in each cell. For each TR, there may be ChIP-seq datasets from different tissues or cell lines, we deduplicate the TR score matrix and keep the TRs with the largest score as the best-matched tissues or dataset for this cell. This generated a normalized TR activity score-by-cell matrix and can be further used to perform clustering, lineage tracing, and other downstream analyses (Fig. S3-S4). After identifying the best-matched ChIP-seq dataset for a TR, we can combine the ChIP-seq peaks with scATAC-seq peaks, and apply the regulatory potential (RP) model (10, 23, 25) to quantitatively evaluate the TR enrichment on its target genes for each cell (Fig. S5). The RP scores reflect the TR regulation ability of its target genes and can be used to construct singlecell GRNs for that TR. Overall, SCRIP will output the TR activity, candidate TR targets, and TR GRNs at single-cell resolution.

### TR Activity Performance Evaluation using PBMC Multiome dataset

To systematically evaluate the performance of SCRIP, we applied it to a peripheral blood mononuclear cell (PBMC) dataset that was produced using the 10X Genomics Multiome platform, which generates scRNA-seq and scATAC-seq in the same cell. We annotated the dataset with cell-type markers from the scRNA-seq dataset and transferred the cell-type labels to the scATAC-seq dataset (Fig. S6a-e). SCRIP successfully finds the key TRs in the corresponding cell types, for example, CEBPA and CEBPB are enriched in monocytes, and PAX5 and BCL6 are enriched in B cells (Fig. 2a, Fig. S7). Also, the cells can be well clustered to their cell type lineages using TR activity alone (Fig. S3). We also compared the consistency of the clustering results with scRNA-seq transferred labels and benchmarked them using normalized mutation information (NMI) and Adjusted Rand index (ARI). Interestingly, clustering using SCRIP TR activity scores shows better consistency with scRNA-seq transferred cell-type compared to existing motif-based methods such as chromVAR, and peak or bin-based methods such as SCALE(11), Signac (8), CisTopic(37), ArchR(38), and SnapATAC(39) (Fig. 2b, Table S1). This result suggests that SCRIP could accurately predict TR activity at the single-cell level, which should show superior performance in determining cell-type lineages.

Next, we compared TR activity with its gene expression. We compared the Spearman correlation coefficients (SCC) distribution of TRs activity scores and its gene expression for both SCRIP and chromVAR. Interestingly, SCRIP has a larger dynamic range for both positive and negative correlations (Fig. 2c, S8a). The majority of chromVAR correlations were around 0, indicating that the motif information might not be able to capture real TR activity. Also, SCRIP identifies more high-confidence TRs for both positive and negative regulators (Fig. S8b). Compared to chromVAR, SCRIP also has generally higher correlations with gene expression in individual cell types (Fig. S8c-k). Also, SCRIP correctly estimated the activity of factors with similar motifs. For example, previous studies have suggested that BCL11A is required for the generation of B progenitor cells (40), while BCL11B activates the transcription of interleukin-2 during T cell activation (41). These two factors share similar motifs but are expressed in distinct lineages (Fig. 2d, Fig. S6c). Consistently, SCRIP predicts BCL11A to be enriched in B-cells and myeloid lineages, while BCL11B is enriched in T and NK-cells (Fig. 2e-f). On the contrary, the chromVAR score shows no significant difference between BCL11A and BCL11B, which is biased by the similar motif sequences (Fig. 2e-f). These results suggest that SCRIP can identify tissue-specific regulations even for factors with similar motifs, which cannot be achieved by motif-based methods.

Finally, we evaluated whether SCRIP can identify cell-type-specific regulations for the same TR. Due to the relative sparsity of TR ChIP-seq datasets, there are slightly more TRs that were only covered by motifs than ChIP-seq (Fig. S3a). We next compared the performance of TR ChIP-seq datasets and motif datasets on the human PBMC dataset. For each TR, we calculated the percentage of cells that selected the motif dataset as the best-matched dataset. Interestingly, for the 335 shared TRs, most of them tend to find ChIP-seq datasets than motif datasets (Fig. S3b). These results suggest although the motifs could serve as a complementary reference to fulfill the TR reference, the ChIP-seq dataset still carries more information than motif datasets for the TRs with both ChIP and motif information. POLR2A is the TR with the most abundant ChIP-seq data in various tissue types (Fig. S1c-d). We tested the ability of SCRIP to find the correct POLR2A ChIP-seq dataset for different single cells. As we expected, for most of the T cells, B cells, and monocytes in PBMC scATAC-seq datasets, SCRIP could successfully identify the corresponding TR ChIP-seq datasets (Fig. S9a-c). If we focused on specific datasets, they were also assigned to the cells with matched cell types (Fig. S9d-f). These results suggest that SCRIP could accurately find the TR datasets with matched cell type information for each cell. In summary, our analyses suggest that SCRIP could accurately predict TR activity at the singlecell level, identify tissue-specific regulations, and find the correct TR dataset for different singlecells.

### TR Targets Evaluation using PBMC scCUT&Tag-pro datasets

The main purpose of ChIP-seq experiments is to find target genes for TR, which is crucial for constructing GRNs. As SCRIP can correctly match the TR ChIP-seq dataset for single cells from different lineages, we asked whether integrating bulk ChIP-seq data and single-cell accessibility could impute the ChIP-seq signals and further identify TR targets at the single-cell level. We thus predicted the TR peaks of each cell with the best match bulk ChIP-seq dataset and applied a modified RP model to infer the putative targets of TR on each cell (Fig. 3a, see Methods). Several single-cell ChIP-seq profiles are available for HMs and a few TRs using scCUT&Tag (20, 22) and scCUT&Tag-pro. While the TR scCUT&Tag data is of low quality, we benchmarked our method using several published HM scCUT&Tag-pro datasets (21).

H3K27ac modification is an active enhancer marker that has been profiled using scCUT&Tag-pro in PBMC(21). We built a reference dataset with active histone modifications including H3K27ac and imputed the H3K27ac signal using the PBMC scATAC-seq dataset (Fig. S2). While the scATAC-seq cells and scCUT&Tag-pro cells are not from the same populations, we cannot compare the performance at the single-cell level. However, when we piled up the H3K27ac scCUT&Tag-pro signal and the SCRIP imputed signal for different cell types, we found that SCRIP could accurately identify the T-cell-specific peaks around STAT4, a TF that plays an important role in T cells (Fig. 3b). Many regions with only scATAC-seq peaks were removed from SCRIP. The CRAMP1 and FAM22A loci show both scATAC-seq signals for monocytes and T cells. However, these two loci do not have H3K27ac signals from the bulk data and they are not output by SCRIP, which were also not observed in the scCUT&Tag-pro data. For the locus of SCL24A42, SCRIP could accurately predict the T-cell-specific H3K27ac signal based on the combination of bulk H3K27ac signal and scATAC-seq peak (Fig. S10). Besides, when we calculated correlations between the real scCUT&Tag-pro RP, SCRIP imputed RP, scATAC-seq RP, and bulk H3K27ac RP, the SCRIP imputed RP shows the highest consistency with the scCUT&Tag-pro RP, indicating its better ability and accuracy in identifying H3K27ac regulated genes (Fig. 3c). More specifically, when comparing the top 1,000 H3K27ac regulated genes in T-cells and monocytes, SCRIP imputed RP could identify more common target genes in scCUT&Tag-pro data than using bulk H3K27ac data directly (Fig. 3d). These results collectively suggest that integrating scATAC-seq data with bulk TR or HM ChIP-seq data could accurately identify their target genes.

### SCRIP Underlies Differentiation Paths for Human HSC Differentiation

TRs are often the driving source of cellular differentiation. To prove that SCRIP can infer TR activity in a complex system and could be potentially used to track cell differentiation, we applied SCRIP on a human hematopoietic stem cell (HSC) differentiation scATAC-seq dataset (42). The HSC differentiation is a well-characterized system, with HSCs differentiating into three different major lineages (Fig. 4a). SCRIP also achieved the second-best result in all methods and shows the best performance in the TR-based method in the clustering performance (Fig. S11e). After identifying TR activity in different HSC subpopulations, we performed a pseudotime analysis and reconstructed the differentiation trajectory of HSC using TR activity (Fig. 4b and Fig. S11a-c, e). The diffusion map of SCRIP suggests that HSC was differentiated into three major directions, CLP (Common Lymphoid Progenitor), Monocytes, and MEP (Megakaryocyte-Erythroid Progenitor), with a little spike towards pDC (Plasmacytoid Dendritic Cells) (Fig. 4b). These directions are perfectly aligned with the known differentiation path of HSCs. By contrast, the diffusion map generated using the original peak count matrix showed a relatively vague separation for different lineages (Fig. S11a). In addition, we have calculated the averaged distance of terminally differentiated cells (monocytes, MEP, CLP, and pDC) versus HSC, SCRIP showed the largest distance compared to using peak-count matrix and chromVAR results, indicating a better lineage separation result (Fig. S11a-d).

Next, we sought to identify the driven TRs for the three major differentiation lineages. We use the average TR activity of CLP, monocytes, and MEP to denote the lineage lymphoid, myeloid and erythroid respectively. Our results correctly distinguish and locate the key TRs into different lineages (Fig. 4c). For example, GATA1 and SPI1 are well-known mutually inhibiting TFs acting as fate-determining regulators in the hematopoietic system. GATA1 specifies the erythroid lineages while SPI1 specifies the myeloid lineage (43, 44), which is highly consistent with the SCRIP results (Fig. 4c). We also found other well-known regulators show high activity in their corresponding lineages, such as HOXA9 for HSCs, CEBPB for myeloid lineages, and TCF4 for lymphoid lineages (Fig. 4d, Fig. S11f-i). Besides, the dynamic changes in the TRs’ activity of the differential lineages indicate their potential role in lineage differentiation (Fig. S11j-y). These results prove that SCRIP enables the trajectory analyses of scATAC-seq with known driver TR activity.

### SCRIP Constructs GRNs in Human Fetal Organ Development

To prove the ability that SCRIP can be applied to diverse tissue types and infer the target genes of TRs, we applied SCRIP to a scATAC-seq dataset of human fetal organs that covers 14 different tissues (45). The TR activity score showed a better performance in clustering compared to the motif-based method chromVAR in almost all these tissues (Fig. 5a, Table S1). To check whether SCRIP could identify the TRs that are involved in the production and maintenance of specific cell types in different organs, we focused on the lung, intestine, and liver datasets. Again, SCRIP could correctly identify the cell-type-specific TRs in these three different organs (Fig. 5b-d, Fig. S12a-c, Table S2). For example, GRHL2 and its downstream direct target gene NKX2-1 form a positive feedback loop to connect lung epithelial cell identity, migration, and lung morphogenesis (46) (Fig. 5b, Bronchiolar and Alveolar Epithelial Cells, BAEC). GATA6 regulates the development of primitive intestinal cells (47) (Fig. 5c, Intestinal epithelial cells, IEC). HNF1A, HNF4A, and GATA4 are well-known hepatocyte TFs in liver tissues (48) (Fig. 5d, Hepatoblasts, HB). These results proved that SCRIP can not only cluster the same cell type with TRs activity but also identify crucial TRs in different cell types using chromatin accessibility data.

Master TRs and their cofactors regulate each other or co-regulate downstream target genes, forming a potential GRN that could modulate cell fate and identities. To validate the ability of SCRIP to establish cell-type-specific GRNs, we inferred the potential target genes of TRs and built the cell-type-specific GRNs for different organs (Fig. 5e-h, Table S3). In the lung, we identified the target genes of GATA3, which is mainly enriched in the lymphoid cells (LC) (Fig. 5b). The target genes of GATA3 mainly contribute to immune functions through responding to interleukin-7 (IL-7) and negatively regulating the differentiation of myeloid cells, which is in line with previous studies (49) (Fig. 5e). In the intestine, MYOD1 controls the differentiation of smooth muscle cells (SMC) by regulating its downstream genes and function (50) (Fig. 5c, f). Finally, we built a co-regulatory GRN of GATA4 and its downstream targets EPAS1(51) in liver hepatoblasts (HB) (Fig. 5g). Although their downstream target genes show a great difference (Fig. S12d-e), the GO analysis suggests that the functions are both enriched in the biosynthetic process. In addition, GATA4 tends to regulate alcohol metabolism, while EPAS1 targets are enriched in response to hypoxia (52, 53) (Fig. 5h). These results show that SCRIP allows identifying the targets of different TRs in diverse cell types and constructing GRNs of multiple TRs in the same cell.

### Disease-Specific GRNs Identified by SCRIP in the Tumor Microenvironment

The target genes of TR can be changed due to different cooperation of co-regulators, especially under disease status (54). We applied SCRIP to a Basal Cell Carcinoma (BCC) tumor microenvironment (TME) dataset (55) to investigate how TRs and their target genes were changed in different cell states under disease status. First, we confirmed the TRs activity is accurately predicted in the corresponding cell types (Fig. 6a). For instance, CEBPG, a TF that promotes cancer development by enhancing the PI3K-Akt signaling pathway (56), was found to be robustly more active in tumor cells than in other cells. In addition, the activity of TFs such as EOMES and TBX21 were higher in immune cells than in tumor cells (Fig. 6a), which is consistent with the role of these TFs in driving lymphocyte differentiation (57, 58).

T cells are the major cytotoxic cells responsible for anti-tumor immunity. Diverse T cell differentiation paths and phenotypes drive the immune response in TME. We performed the pseudo-time analysis of T cells in TME using the TR activity from SCRIP, which uncovered two distinct paths. The first differentiation path is from naïve CD4 T cells to T follicular helper (Tfh) cells, for which IRF4 is gradually activated in Tfhs (Fig. 6b). The IRF4 activity was significantly increased in Tfh, and the function of its target genes was also enriched in lymphocyte activation and differentiation (Fig. S13a). These analyses are consistent with the IRF4 function in Tfh cell expansion (59). Another path is from naïve CD8 T cells to terminal T exhaustion (TEx) cells. BATF, a key regulator of T cell exhaustion (60), has higher activity in terminal TEx (Fig. 6c). Consistently, the BATF target genes tend to have an immunosuppressive effect on terminal TEx cells (Fig. S13b). These analyses suggest that the TR activity and targets inferred by SCRIP could be used to track cell state changes under the disease condition.

Interestingly, we found that the activity of JUNB are both higher in naïve CD8 T and terminal TEx (Fig. S13c). We then checked the target genes of JUNB between these two cell types. Although most targets were shared, there are a considerable number of differential targets between these two stages (Fig. 6d, e). We asked whether JUNB has different functions in naïve CD8 T cells and terminal TEx cells, then we examined their differential target genes and built cell-type-specific GRNs for JUNB (Fig. S13d, Fig. 6f, g, Fig. S14). We found that PRMT5, which is critical for the transition of naïve T cells to the effector or memory phenotype (61), is presented in the JUNB GRNs only in naïve CD8 T cells (Fig. 6e, f). In contrast, CTLA4, which could encode a protein that transmits an inhibitory signal to T cells and its upregulation has been described as a marker of T cell exhaustion in chronic infections and cancer (62, 63), has a high RP score in JUNB GRNs in terminal TEx (Fig. 6e, g). The function enrichment results suggest that JUNB mainly tends to function as a positive regulator of T cell activation and migration to lymphoid organs, while negatively modulating the immune system process in terminal TEx cells. In summary, our analyses suggest that SCRIP can identify cell-type-specific GRNs as well as uncover disease-specific GRNs in complex biological systems.

## Discussion

In this study, we present SCRIP, a computational workflow for single-cell gene regulation inference by large-scale data integration. We first built a manually curated and comprehensive epigenome reference dataset including 11k human and 9k mouse TR ChIP-seq data. Based on the reference, we developed a method that can evaluate TR activity and build GRNs at the single-cell resolution using scATAC-seq. Our method achieves better performance compared to the previous motif-based methods in terms of clustering accuracy, consistency with gene expression, and ability to discriminate factors within the same family. We applied SCRIP to four different biological systems, including PBMC, HSC differentiation, human fetal organ development, and the BCC tumor microenvironment. SCRIP does not merely identify the key TRs in different cell types under diverse biological settings. In addition, the TR activity predicted by SCRIP could be used to trace the cell lineages and identify lineage-specific regulators. The single-cell GRNs constructed by SCRIP enable the identification of the co-regulation relationship between different TRs and reveal the disease-associated GRNs in the terminal exhausted T-cells from the tumor microenvironment.

Although in SCRIP, the ChIP-seq-based method outperforms motif-based methods in many aspects, there are still several limitations. First, our method significantly relies on the data quality of the ChIP-seq datasets. After filtering the 20k human and mouse datasets, there are only 4k ChIP-seq datasets with good data quality. This significantly reduced the number of TRs as well as different types of tissues covered by our reference. To compensate for this, we also integrate the motif scanning results into the TR reference. Second, there might be potential batch effects between the TR ChIP-seq data with the scATAC-seq data. To avoid this, for each cell we score TR enrichment using multiple TR ChIP-seq from different tissues, and only keep the one with the highest TR enrichment score, which is usually from the same cell type. This could partially solve the batch effect for TRs with a large number of datasets, but may not be appropriate for TRs with few numbers of matched datasets. Third, the experiment of the public TR ChIP-seq may have been performed with different perturbations, which may alter the TR’s binding sites and introduce biases to our results. Finally, the bulk-level ChIP-seq datasets have the probability of losing the signals on rare populations, which also impacts the results of binding sites and target genes for our method, especially for some minority populations.

We foresee several ways to further improve our method. First, there will be an increasing number of TR ChIP-seq, CUT&RUN, and CUT&Tag datasets in the future. The first version of CistromeDB includes 13,366 human and 99,53 mouse epigenome datasets, while the number almost doubled to 25,000 human and 22,000 mouse epigenome datasets after only 2 years (16, 26). Large consortiums like ENCODE, and Epigenome Roadmap will also generate a great number of TR datasets with high quality. With the development of scCUT&RUN and scCUT&Tag-pro, we could also integrate the single-cell TR dataset into our reference for annotating TRs from scATAC-seq in other cell types. These expanded references will improve the performance of our method for predicting TR activity. Second, machine learning algorithms, such as generative adversarial networks (GAN) could be used to generate more TR ChIP-seq datasets *in silico*. Finally, although we demonstrated that SCRIP is powerful in predicting TR activity and GRNs for scATAC-seq, we could potentially extend its applications to scRNA-seq. The CistromeDB also has a decent collection of public ATAC-seq and DNase-seq datasets, which could be used to infer the chromatin accessibility for scRNA-seq, then infer the TR activity using the predicted accessibilities. In addition, gene expression correlation could be considered to increase the accuracy of constructing single-cell GRNs. With the implementation of those features, we anticipate SCRIP to help researchers identify driver TRs and interpret single-cell GRNs in different biological areas.

## Declarations

### Ethics approval and consent to participate

Not applicable

### Consent for publication

Not applicable

### Data availability

PBMC multiome dataset is available on the 10X genomic website (https://www.10xgenomics.com/resources/datasets/pbmc-from-a-healthy-donor-granulocytes-removed-through-cell-sorting-10-k-1-standard-2-0-0). scCUT&Tag-pro H3K27ac dataset was obtained from their original studies: https://zenodo.org/record/5504061. Other datasets analyzed during the current study are available in the GEO with the following accession: HSC (GSE96769), human fetal organ (GSE149683), BCC tumor microenvironment (GSE129785).

### Code availability

SCRIP is an open-source python package with source code freely available at: https://github.com/wanglabtongji/SCRIP.

### Authors’ contributions

C.W. conceived and supervised the project. X.D. designed and implemented the SCRIP algorithm. X.D. collected and preprocessed the ChIP-seq and motif datasets, and built the SCRIP index. X.D., K.T., and Y.X. evaluated the performance. X.D. performed the analysis of PBMC. K.T. performed the analysis of HSC and human fetal organs. Y.X. performed the analysis of T cells in the tumor microenvironment. X.D., K.T., Y.X., and C.W. wrote the manuscript with the help of other authors. All authors read and approved the final manuscript.

