## Supplementary Figures for "Single-cell Gene Regulation Network Inference by Large-scale Data Integration"

**Supplementary Figure S1**

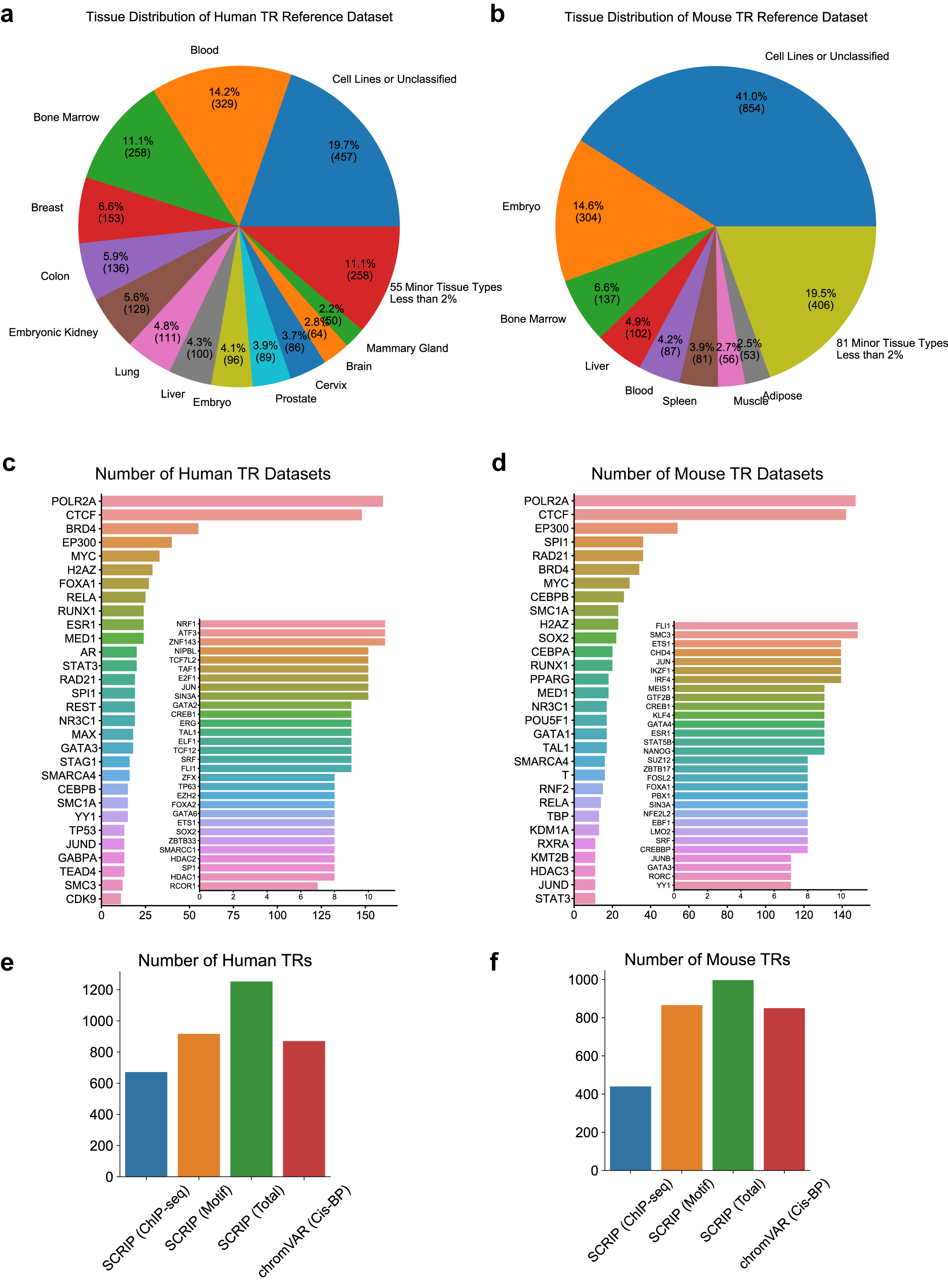

**Overview of TR reference datasets of human and mouse**

**a-b.** Tissue distribution of human and mouse TR reference datasets. Cell lines and minor tissue types are not fully marked.

**c-d.** Histogram of TR of human and mouse reference datasets. Only the top 60 TRs with datasets are shown.

**e-f.** The number of covered TRs of human and mouse reference datasets in SCRIP and chromVAR.

**Supplementary Figure S2**

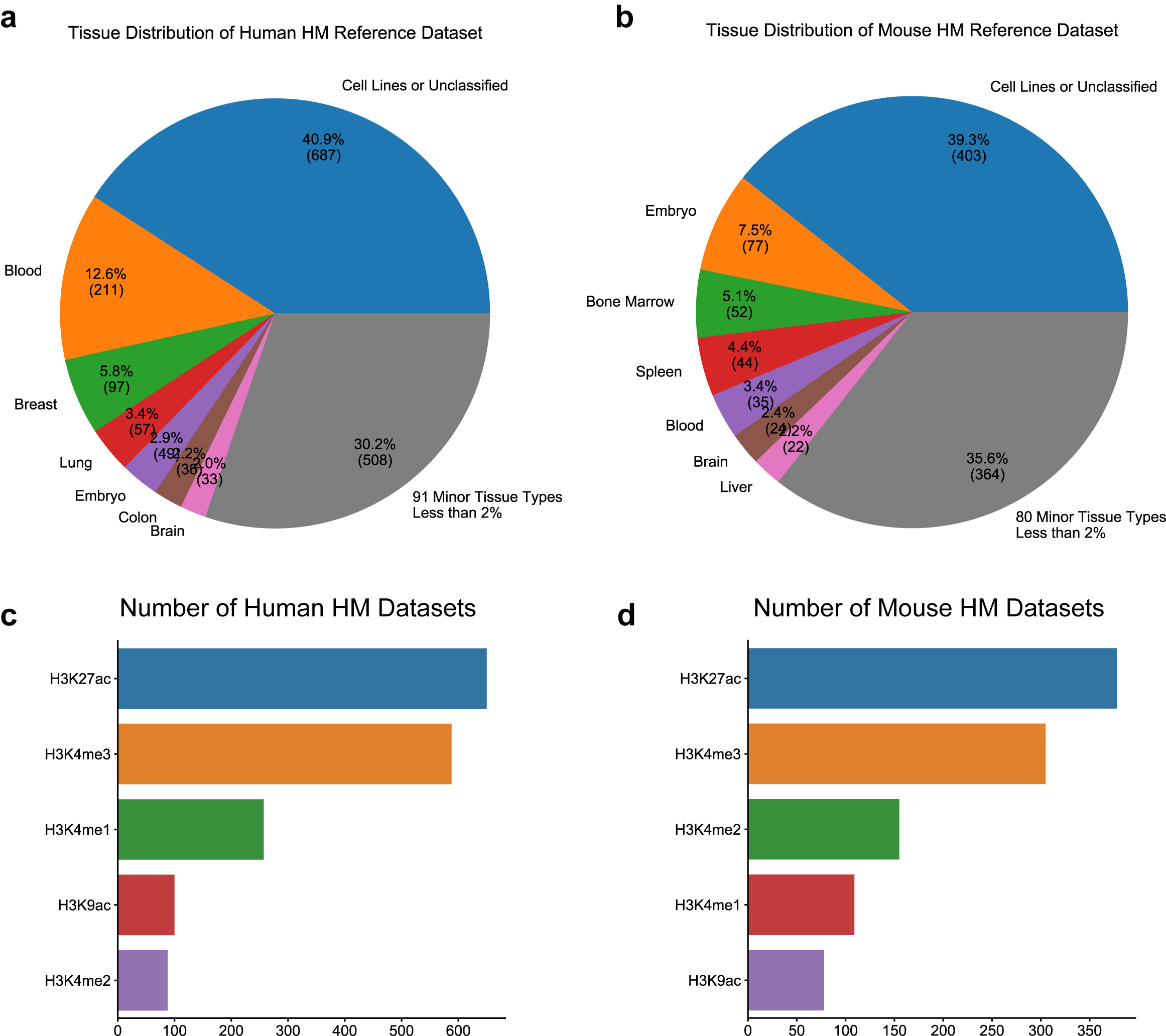

**Overview of histone modifications reference datasets of human and mouse**

**a-b.** Tissue distribution of human and mouse histone modifications reference datasets. Cell lines and minor tissue types are not fully marked.

**c-d.** Histogram of histone modifications of human and mouse reference datasets.

**Supplementary Figure S3**

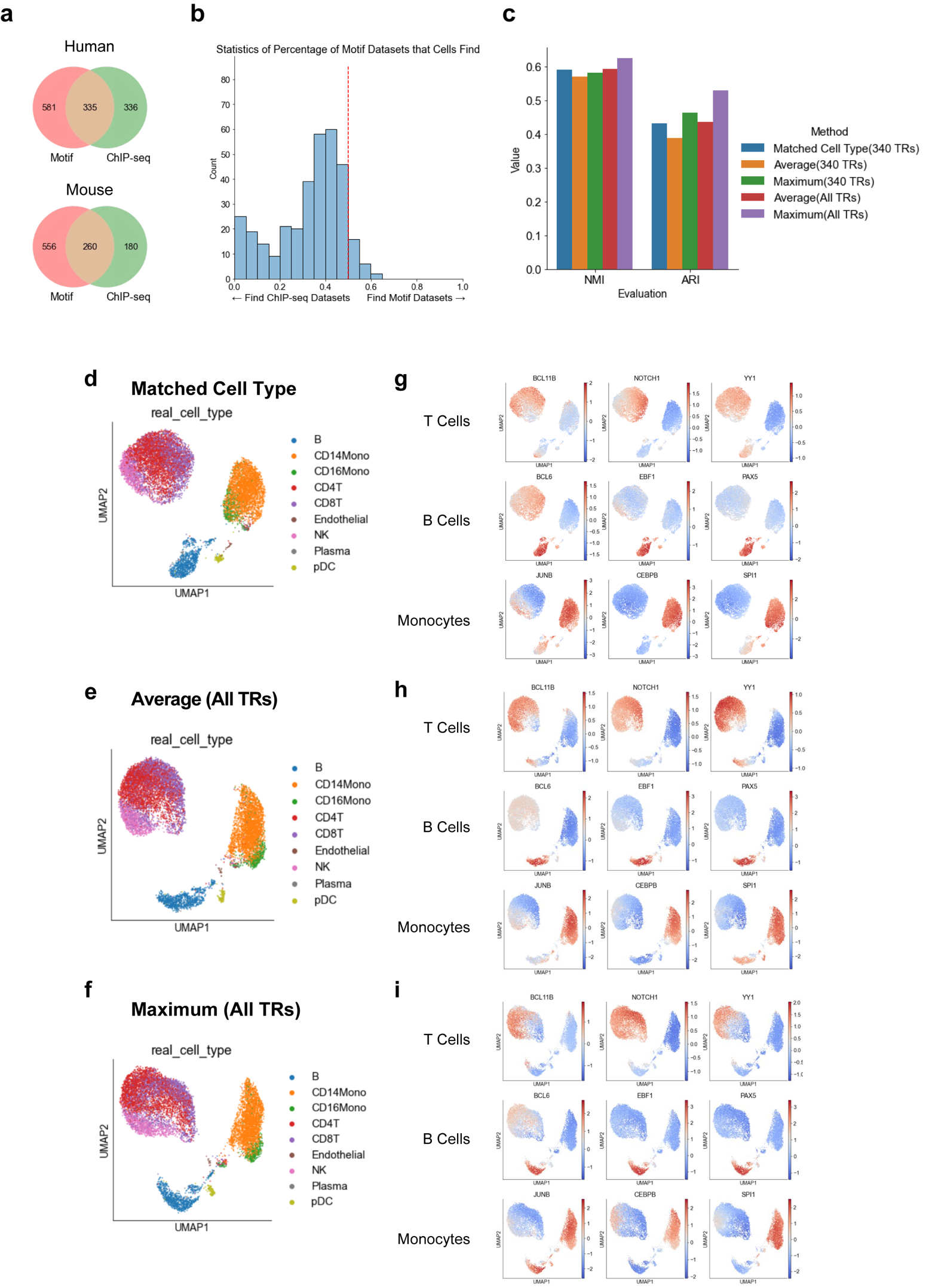

**Comparison of different datasets selection strategies**

**a.** Overlap of TRs from motif datasets or ChIP-seq datasets.

**b.** Distribution of percentage of cells that select motif as best matched dataset. x-axis, percentage of cells that select motif as best matched dataset, y-axis, number of TRs that have this percentage.

**c.** Consistency of scATAC-seq clusters to scRNA-seq transferred cell types for different scATAC-seq tools, accuracies were evaluated using both NMI and ARI.

**d-i.** TR enrichment distributions of different data selection strategies. **d-f.** Clustering results of different data selection strategies, **d.** cell-type-matched strategy, **e.** average strategy, **f.** maximum strategy. **g-i.** TR enrichment distributions of different data selection strategies, **g.** cell-type-matched strategy, **e.** average strategy, **i.** maximum strategy

**Supplementary Figure S4**

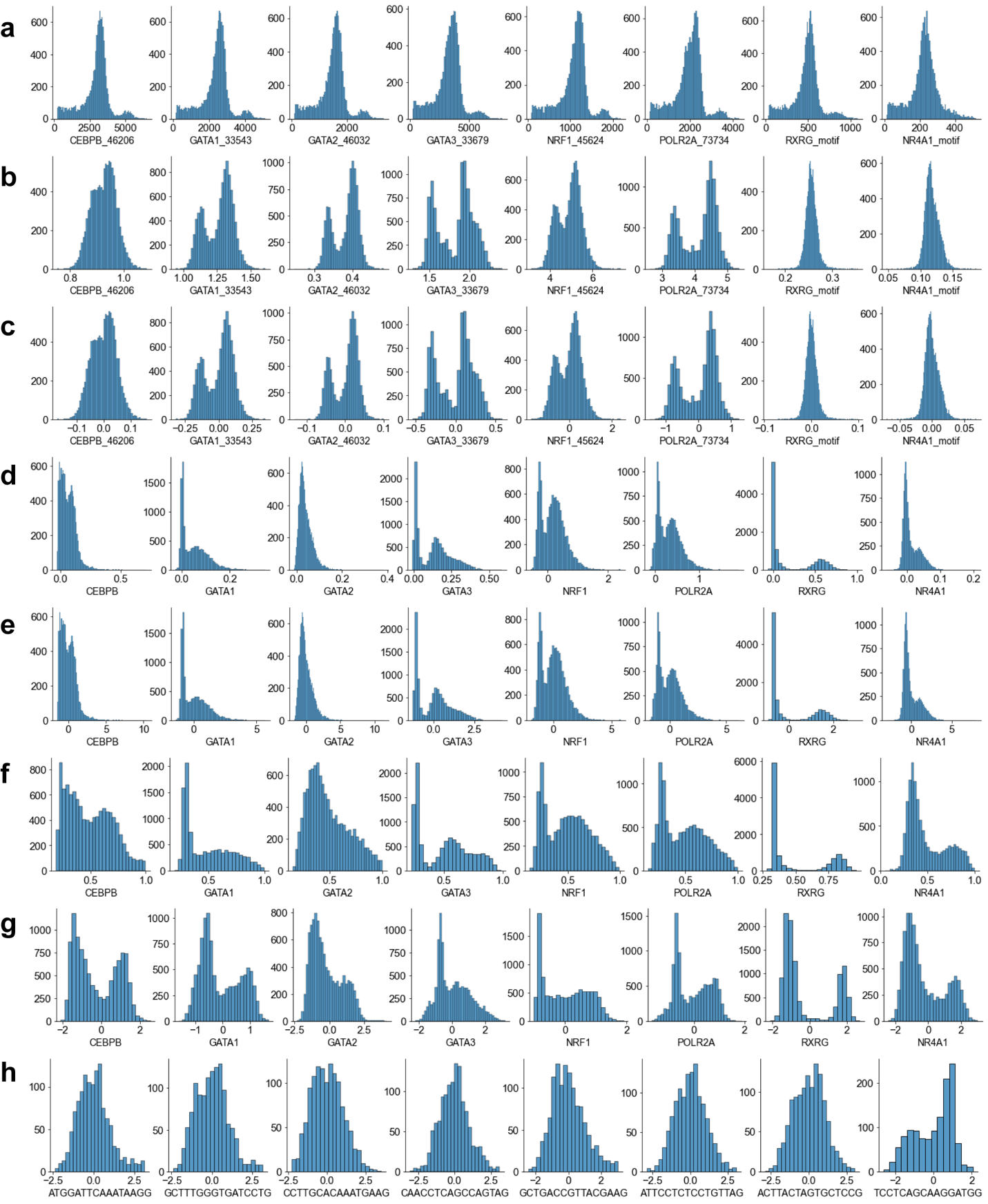

**Distribution changed during the calculation and normalization**

**a-g.** In each figure, the y-axis denotes the number of scores. The plots suggest the distribution in all cells of **a.** the number of original peak overlaps. **b.** the scores after normalizing reference peaks number and scATAC-seq peaks length. **c.** the scores after subtracting the mean of scores of the same dataset. **d.** the scores after summarizing the dataset to TR by the maximum strategy. **e.** the scores after first z-score normalization. **f.** the scores after sigmoid transformation. **g.** the scores after second z-score normalization.

**h.** The plots suggest the distribution of TRs in cells.

**Supplementary Figure S5**

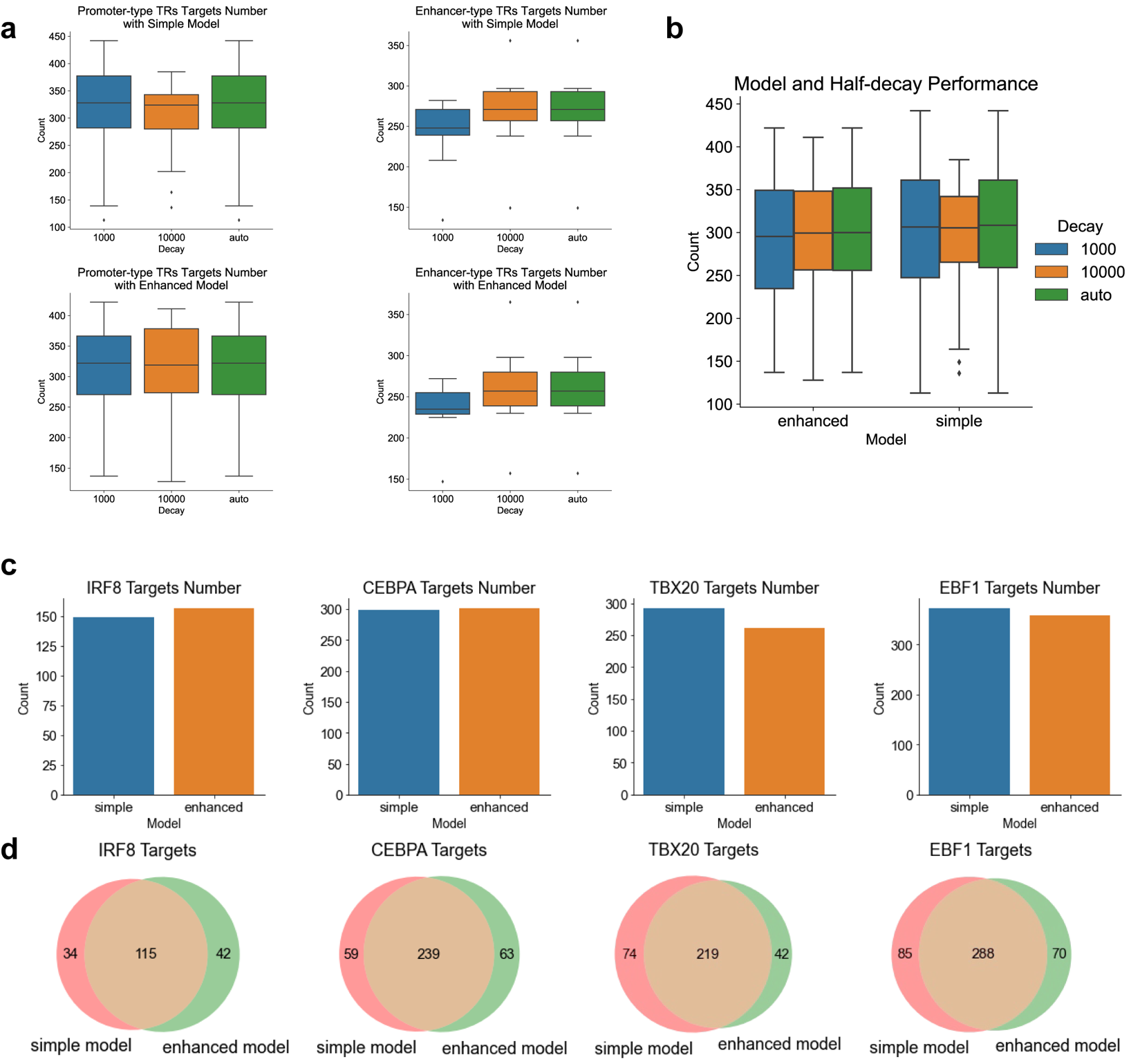

**Evaluation of different RP models on target identification**

**a.** Performance of RP models with different half-decay distances. 1k, all TRs using 1k as half-decay distance; 10k, all TRs using 10k as half-decay distance; auto, automatic determine decay distance based on peaks enrichment on promoter regions.

**b.** Performance of simple and enhanced RP models. The simple model is from Cistrome-GO, and the enhanced model is from MAESTRO.

**c.** Number of true targets among the top 500 targets for different factors.

**d.** Overlaps of true targets for different RP models.

**Supplementary Figure S6**

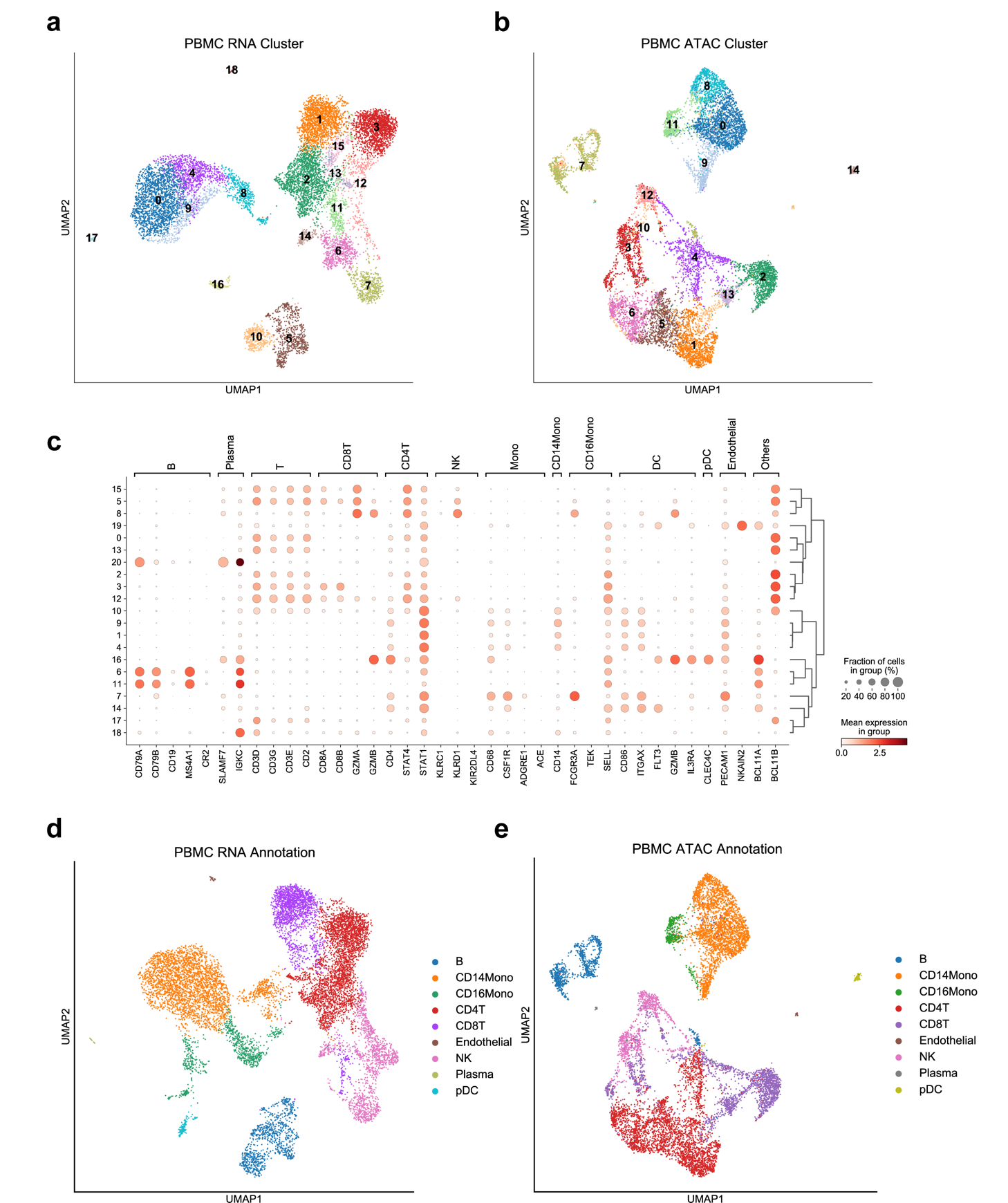

**PBMC scRNA-seq dataset and cell-type annotation**

**a.** UMAP which is based on PBMC scRNA-seq gene expression shows Louvain clustering.

**b.** UMAP which is based on PBMC scATAC-seq peak count shows Louvain clustering.

**c.** Gene markers were used to annotate the PBMC scRNA-seq dataset.

**d.** UMAP which is based on PBMC scRNA-seq gene expression shows the cell-type annotation.

**f.** UMAP which is based on PBMC scATAC-seq peak count shows the cell-type annotations that were transferred from scRNA-seq by barcodes.

**Supplementary Figure S7**

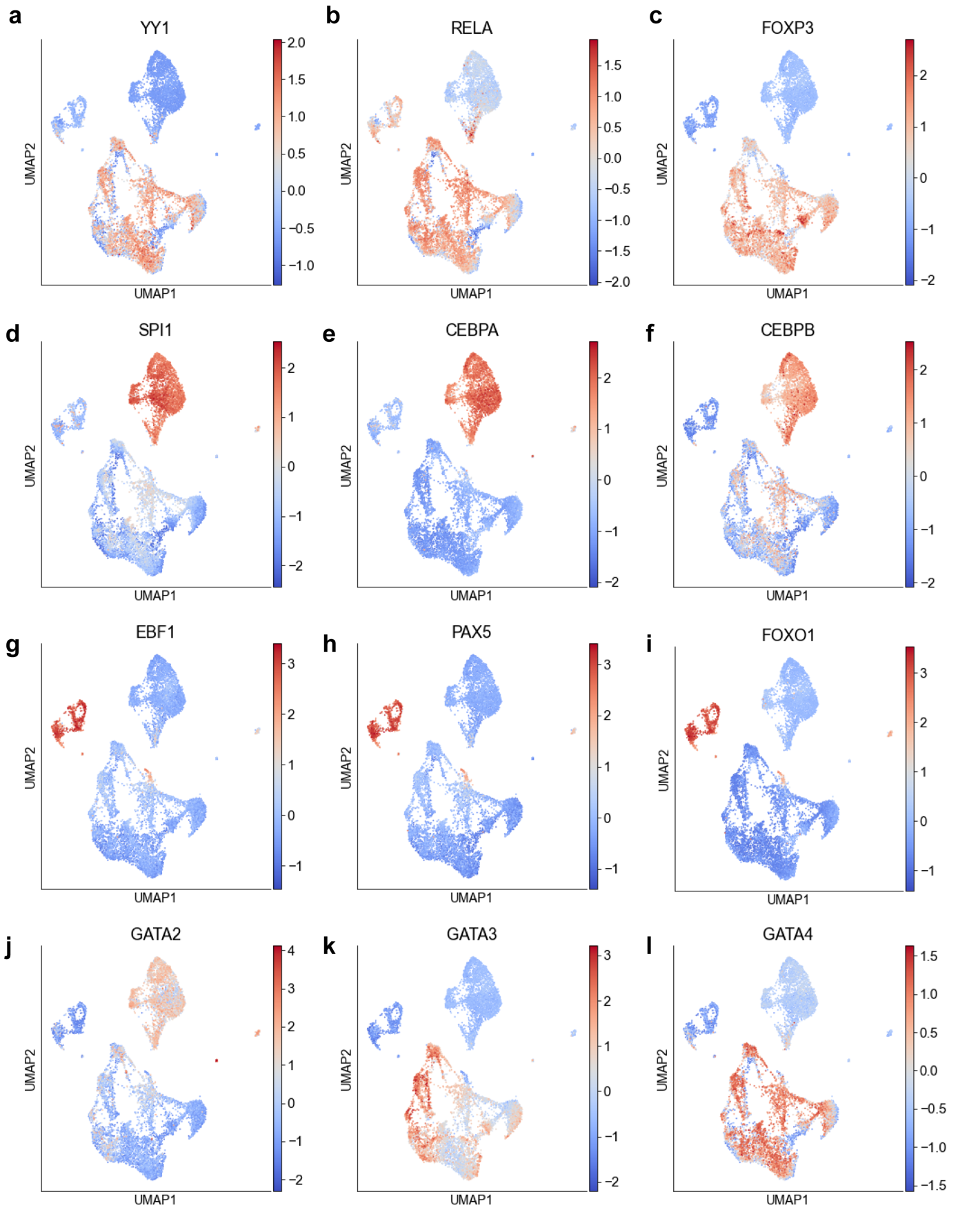

**Well-known TR activity in the diverse cell types of PBMC**

**a-c.** UMAP shows the YY1, RELA, and FOXP3 activity, which are important TRs in T cells.

**d-f.** UMAP shows the SPI1, CEBPA, and CEBPB activity, which are important TRs in monocytes.

**g-i.** UMAP shows the EBF1, PAX5, and FOXO1 activity, which are important TRs in B cells.

**j-l.** UMAP shows the GATA2, GATA3, and GATA4 activity, which share similar motifs (not shown).

**Supplementary Figure S8**

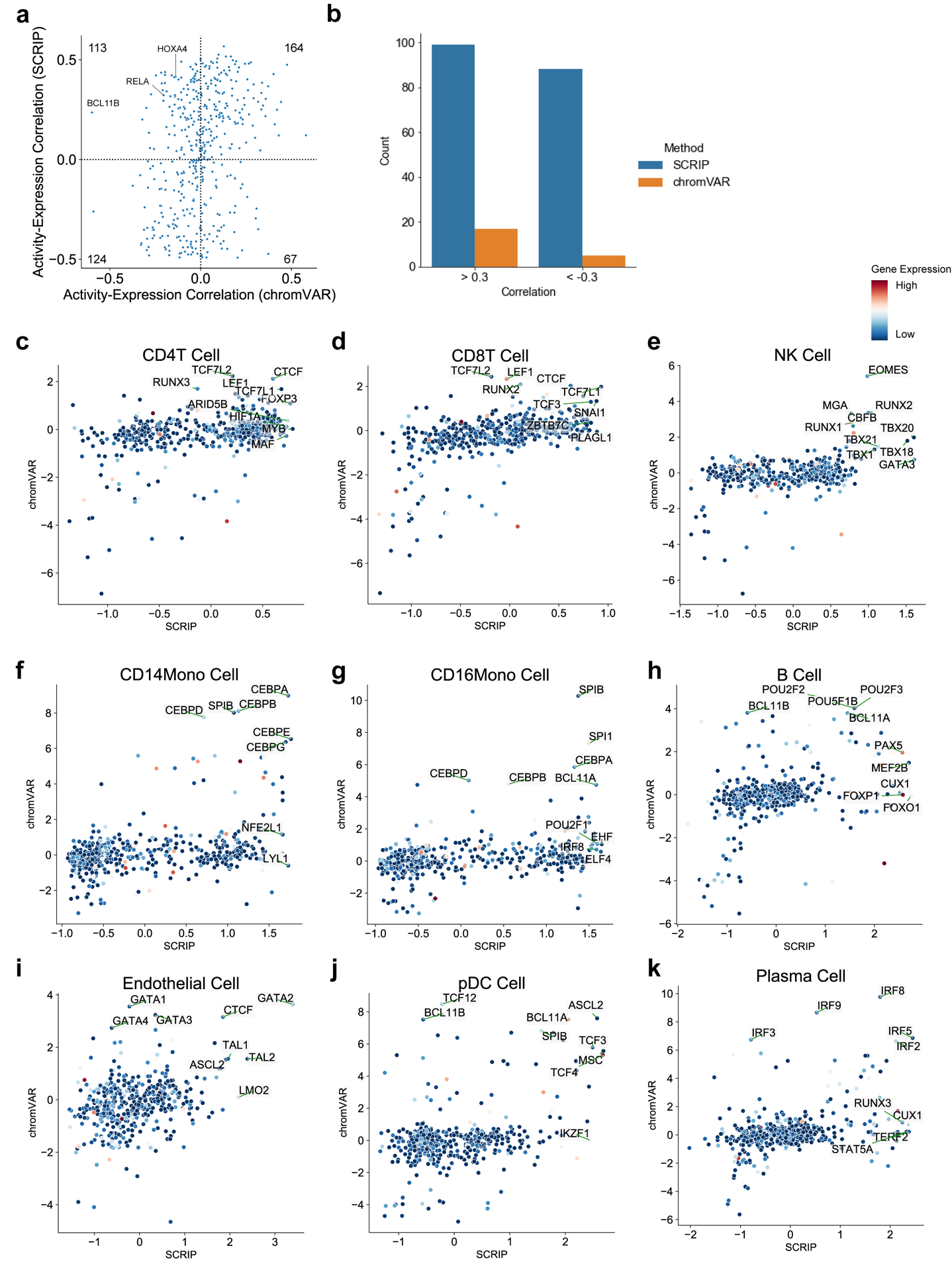

**Comparison of the TR activity with gene expression between SCRIP and chromVAR**

**a.** TR activity and its gene expression spearman correlation. X-axis: correlation of chromVAR TF enrichment z-score and TF expression; Y-axis: correlation of SCRIP TR activity and TR expression. The number of each corner denotes the number of TR in each quadrant.

**b.** Number of High-confident positive regulations and negative regulators for SCRIP and chromVAR. High-confident regulators were defined as SCC > 0.3 and < -0.3, and p-value < 0.01.

**c-k.** Scatter plot of average TR activity of two methods and average gene expression in different cell types of PBMC. TRs in the top 1% either SCRIP or chromVAR were marked. The color of each dot denotes the average gene expression in the cell type.

**Supplementary Figure S9**

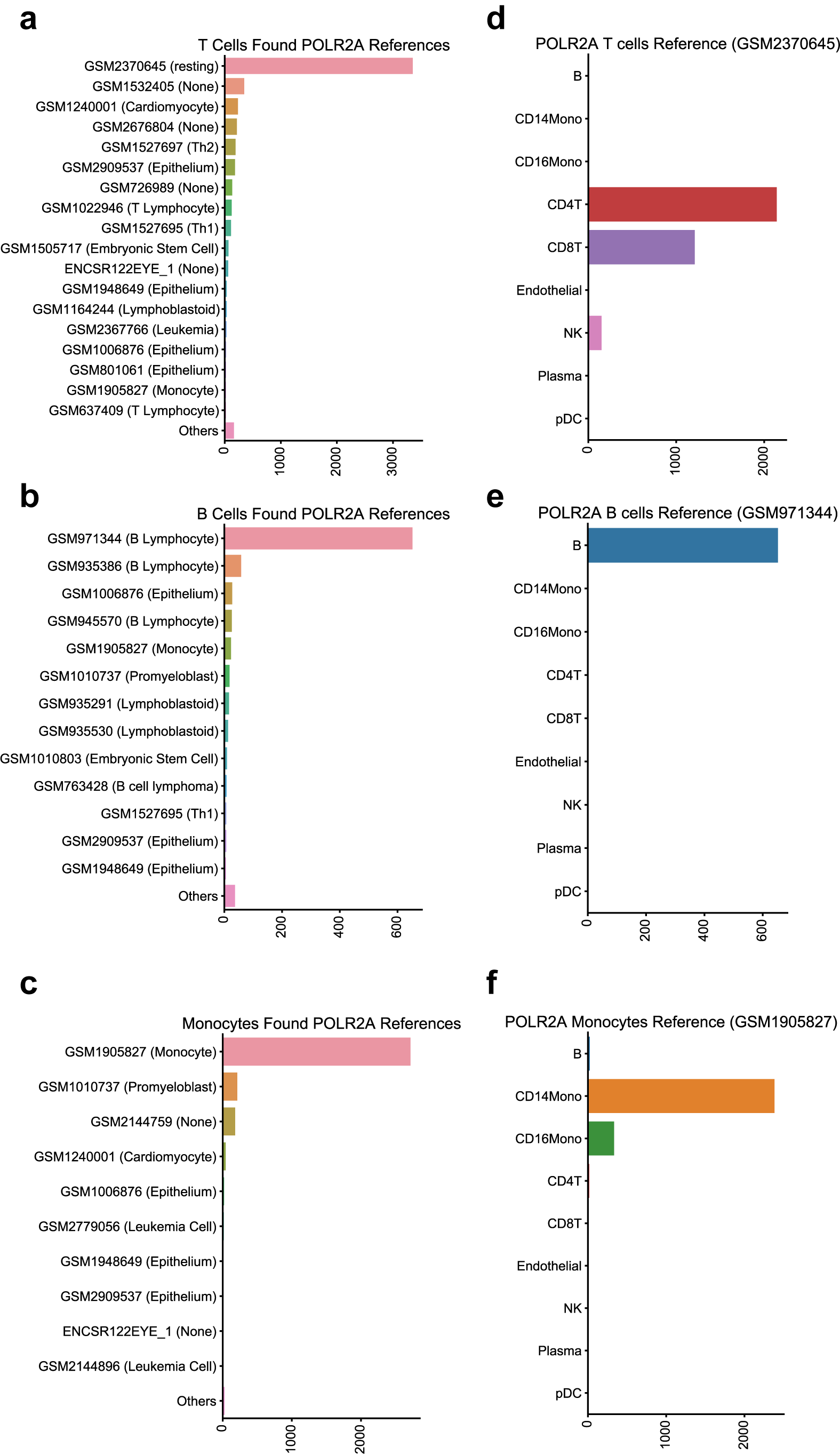

**SCRIP found the matched cell type between single-cell data and bulk data**

**a-c.** Histogram showing the majority of found reference datasets in T cells, B cells, and monocytes.

**d-f.** Histogram to show what cells matched the top 3 found on reference datasets.

**Supplementary Figure S10**

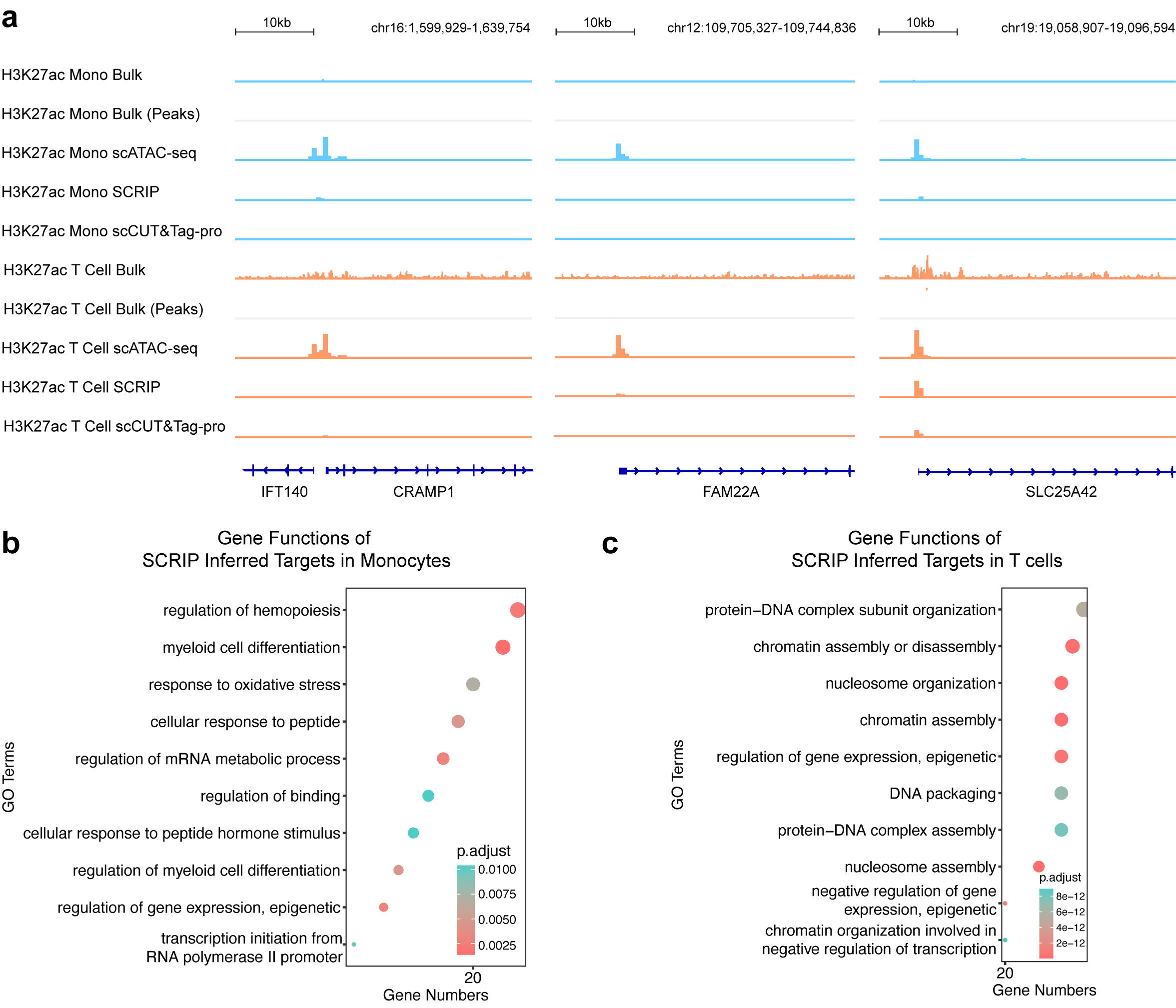

**Genome track of monocytes and T cells on H3K27ac signals at CRAMP1, FAM22A, and SLC25A4 and GO analyses.**

**a.** Light blue: Monocytes; Orange: T cells. Bulk tracks are read level; scCUT&Tag-pro, SCRIP-inferred, and scATAC-seq tracks are 500 bp bin level. In single-cell tracks, the height of the signal denotes the normalized cell number.

**b.** GO results showed the terms enriched of top H3K27ac target genes in monocytes.

**c.** GO results showed the terms enriched of top H3K27ac target genes in T cells.

**Supplementary Figure S11**

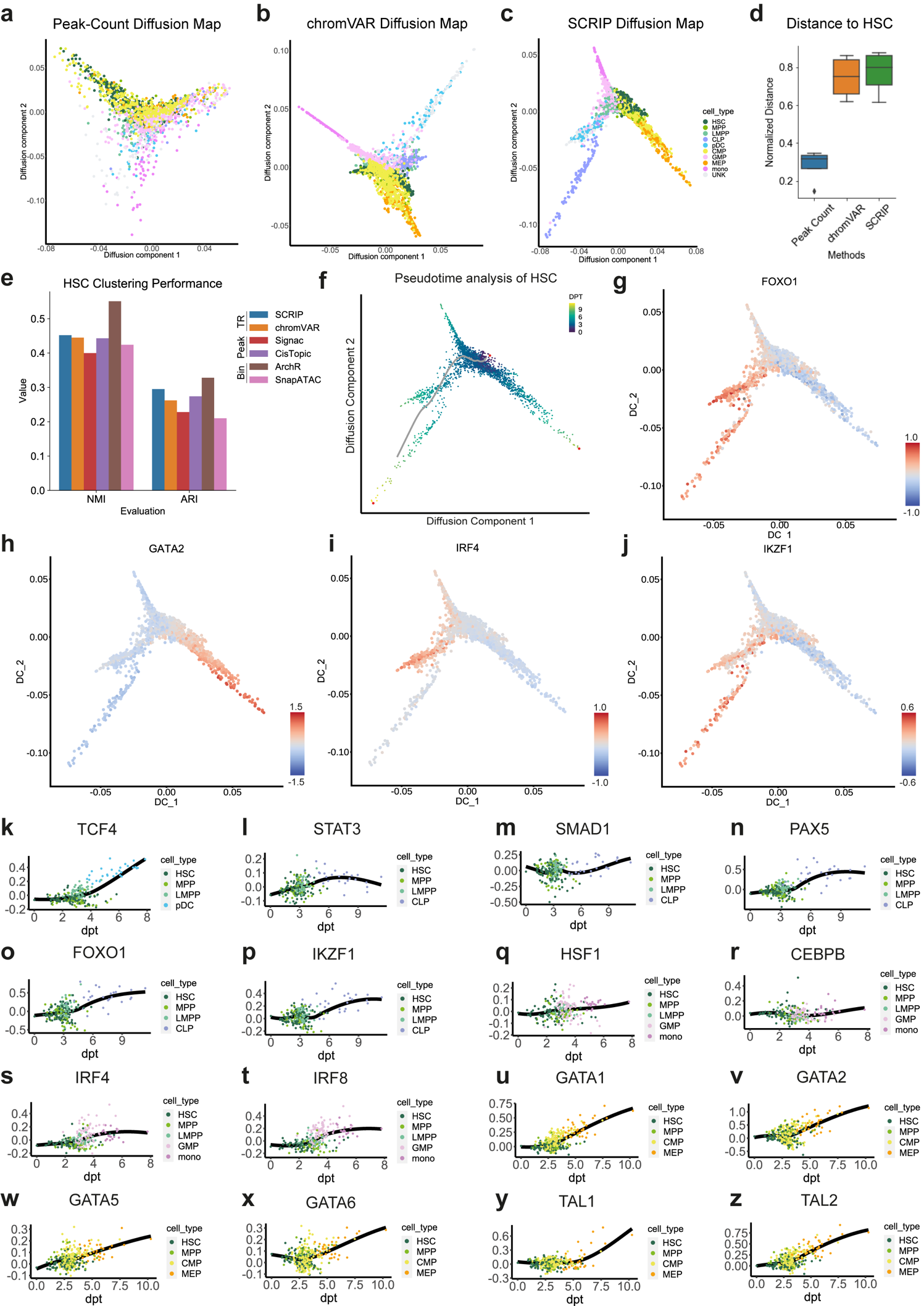

**Pseudo time of HSC and changes of important TR activity during differentiation**

**a-c.** Diffusion map using **a.** original peak counts. **b.** chromVAR z-scores. **c.** SCRIP TR activity scores.

**d.** Normalized distances of 4 terminal cell types to HSC between three methods.

**e.** The clustering performance of HSC among methods. X-axis: NMI and ARI matrix. TR-based methods: SCRIP, chromVAR; Peak-based methods: CisTopic; Bin-based methods: ArchR, SnapATAC. Y-axis: NMI score and ARI scores

**f.** Diffusion map of HSC pseudo-time. Dark purple denotes the early stages, and yellow denotes the late stages.

**g-j.** Projecting FOXO1, GATA2, IRF4, and IKZF1 activity onto the diffusion map.

**k-z.** TR activity changes during cell differentiation. **k.** HSC to pDC; **l-p.** HSC to CLP; **q-t.** HSC to monocytes; **u-z**. HSC to MEP.

**Supplementary Figure S12**

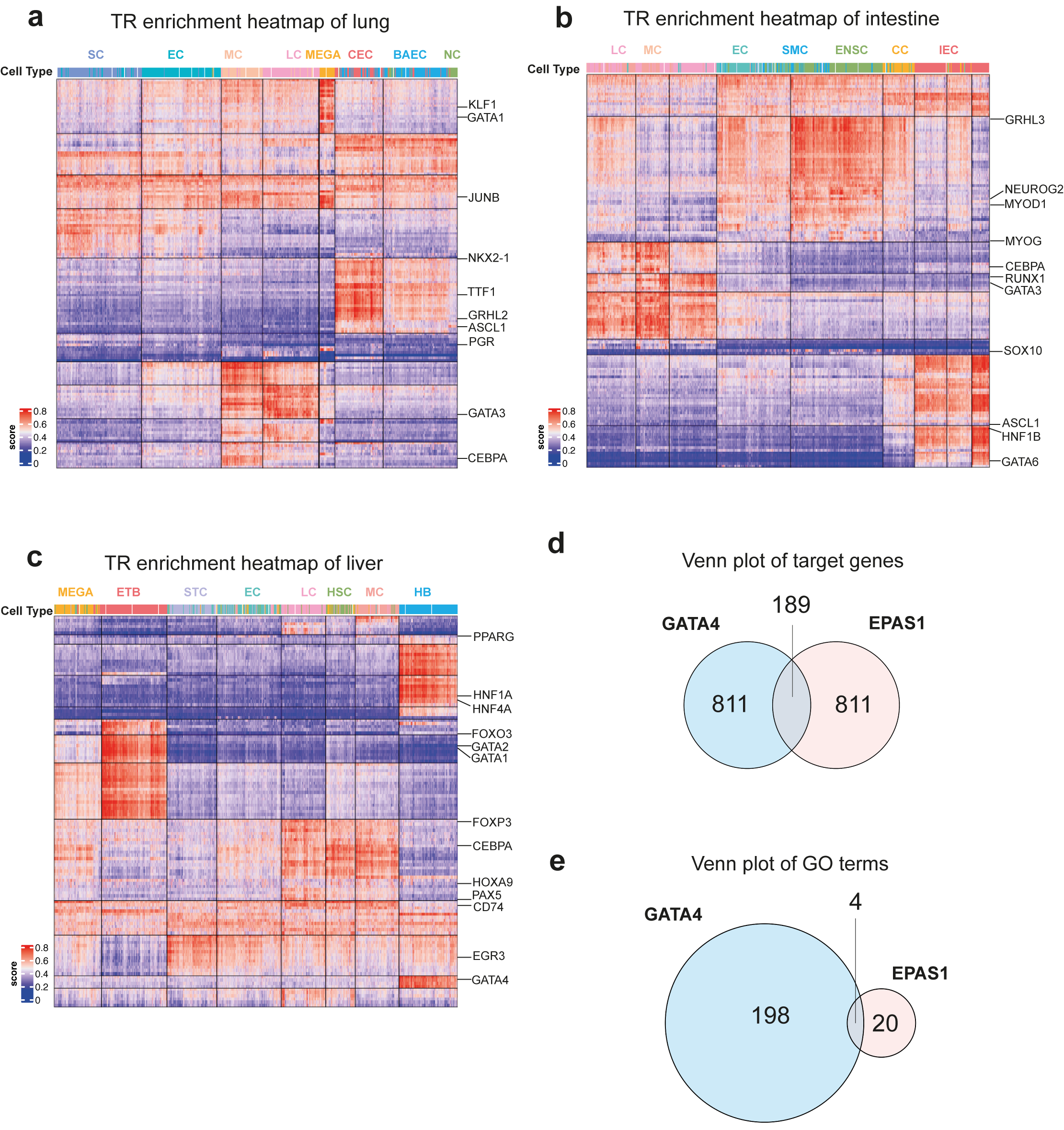

**TR activity clustering of three human organ datasets and differences in co-regulation**

**a-c.** Heatmap of TR activity with different cell types in the lung, intestine, and liver.

**d.** Overlap of top 1,000 target genes of GATA4 and EPAS1 in HB of human liver.

**c.** Overlap of enriched GO terms of GATA4 and EPAS1 target genes in HB of the human liver.

**Supplementary Figure S13**

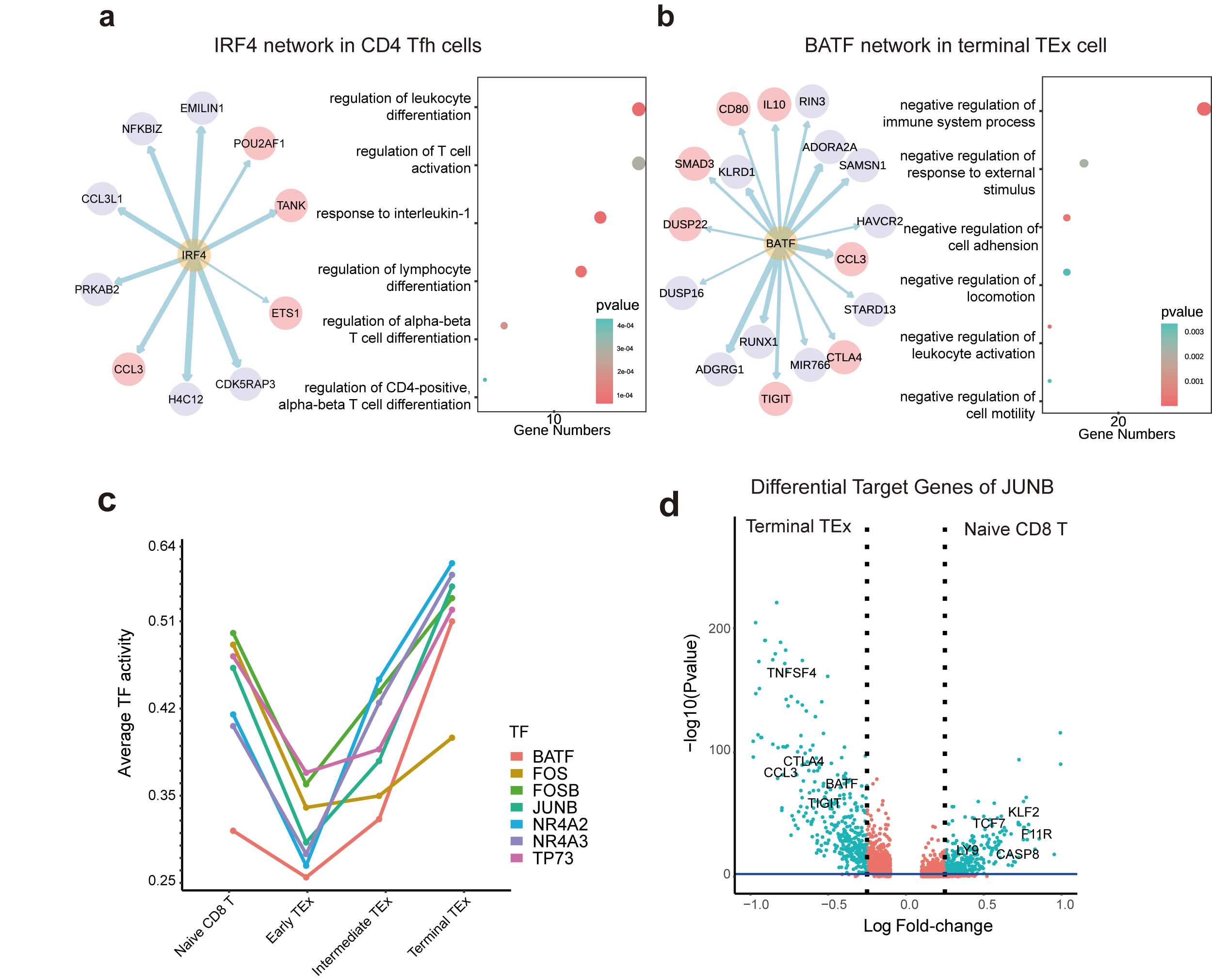

**Key TR GRNs and differential target gene determination**

**a.** (left) Inferred IRF4 GRN in CD4 Tfh cells. Pink circles denote the target genes that are supported by previous studies. (right) GO results showed the terms enriched of IRF4 target genes in CD4 Tfh cells.

**b.** (left) Inferred BATF GRN in terminal TEx cells. Pink circles denote the target genes that are supported by previous studies. (right) GO results showed the terms enriched of BATF target genes in terminal TEx cells.

**c.** Line plot to show the changes of 6 TRs activity during the differentiation from naive CD8 T cells to terminal TEx cells. Y-axis: average TR activity of each cell type.

**d.** Volcano plot to show the different target genes of JUNB between naive CD8 T cells and terminal TEx cells.

**Supplementary Figure S14**

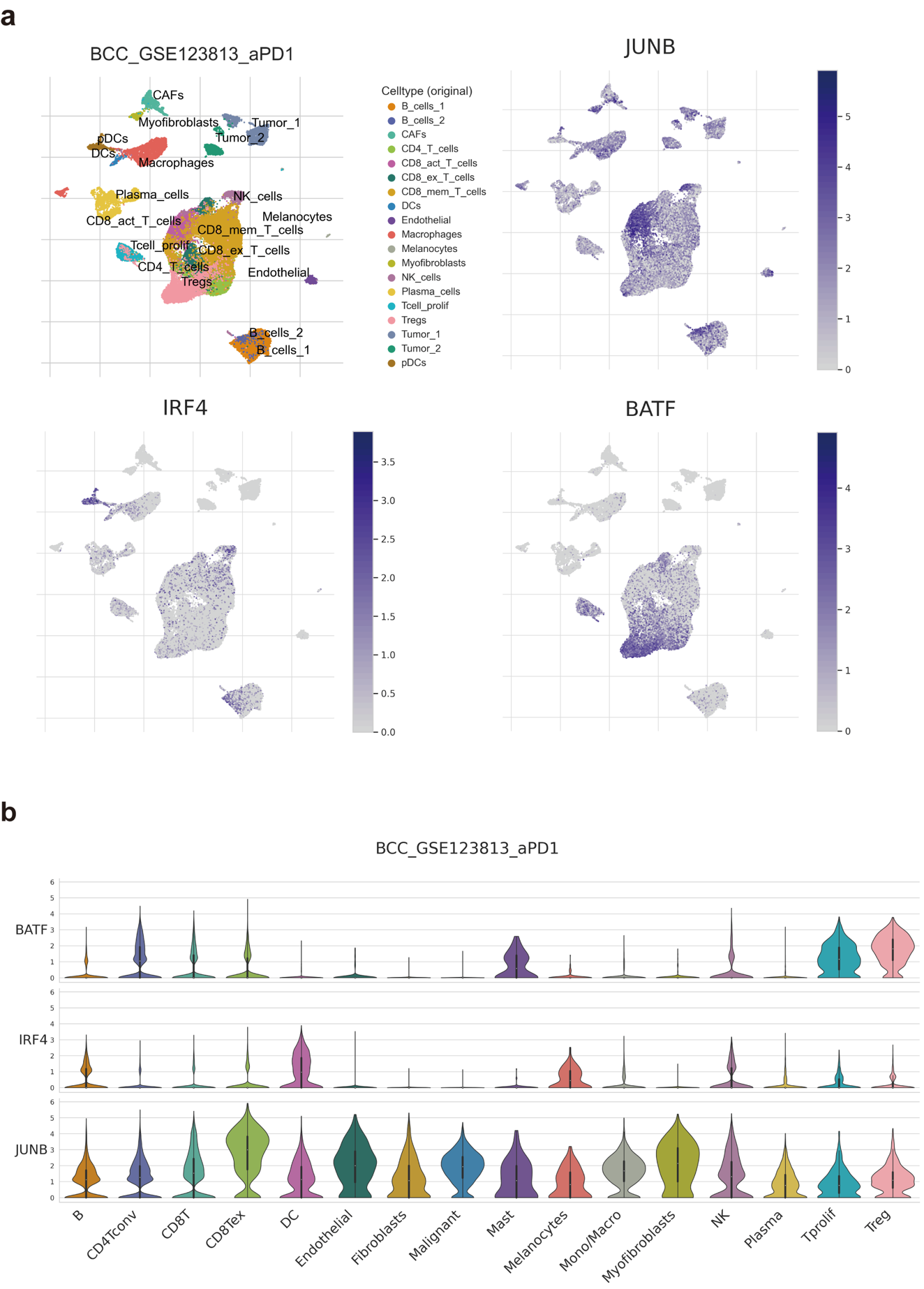

**Gene expression of example TRs in BCC dataset**

**a.** UMAP of cell identities of BCC datasets and the gene expression of JUNB, BATF, and IRF4 obtained from the TISCH database.

**b.** Violin plot to show the gene expression of JUNB, BATF, and IRF4 in different cell types of BCC datasets.

**Supplementary Tables**

**Supplementary Table S1**

Clustering performance of different methods on different tissues. The numbers suggest the NMI and ARI scores. In the table, "x" means that fails to run because of the size of the data, and the blank means that we do not run this tool on the dataset.

**Supplementary Table S2**

Specific-cell-type TRs with literature supporting. The numbers in the brackets denote the PubMed ID of the paper that supports the TR playing roles in this cell type.

**Supplementary Table S3**

TF-target pairs with literature supporting. The numbers in the brackets denote the PubMed ID of the paper that supports the gene that is the target of this TR.
